# The coordinated localization of mRNA to centrosomes facilitates error-free mitosis

**DOI:** 10.1101/2020.04.09.034892

**Authors:** Pearl V. Ryder, Junnan Fang, Dorothy A. Lerit

## Abstract

Centrosomes are microtubule-organizing centers required for error-free mitosis and embryonic development. The microtubule-nucleating activity of centrosomes is conferred by the pericentriolar material (PCM), a composite of numerous proteins subject to cell cycle-dependent oscillations in levels and organization. In diverse cell types, mRNAs localize to centrosomes and may contribute to changes in PCM abundance. Here, we investigate the regulation of mRNA localization to centrosomes in the rapidly cycling *Drosophila melanogaster* embryo. We find that RNA localization to centrosomes is regulated during the cell cycle and developmentally. We identify a novel role for the fragile-X mental retardation protein (FMRP), which localizes to pericentrosomal RNA granules, in the post-transcriptional regulation of centrosomal RNA. Further, the mis-targeting of a model centrosomal mRNA, *centrocortin* (*cen*), is sufficient to alter cognate protein localization to centrosomes and impair spindle morphogenesis and genome stability.

## Introduction

The centrosome is a multi-functional organelle that serves as the primary microtubule-organizing center of most animal cells and comprises a central pair of centrioles surrounded by a proteinaceous matrix of pericentriolar material (PCM) (Conduit, Wainman, & Raff, 2015). During mitosis, centrosomes help organize the bipolar mitotic spindle and function to ensure the fidelity of cell division. In interphase, centrosomes contribute to cell polarization, intracellular trafficking, and ciliogenesis (Vertii, Hehnly, & Doxsey, 2016).

Cell cycle-dependent changes in PCM composition contribute to functional changes in centrosome activity. Upon mitotic entry, centrosomes undergo mitotic maturation, a process by which centrosomes augment their microtubule-nucleating capacity through the recruitment of additional PCM (Palazzo, Vogel, Schnackenberg, Hull, & Wu, 1999). This process is reversed upon mitotic exit by PCM shedding (Magescas, Zonka, & Feldman, 2019; Mittasch et al., 2020). These dynamic oscillations in PCM composition and organization are essential for centrosome function, and their deregulation is associated with developmental disorders, increased genomic instability, and cancer (Conduit et al., 2015; Nigg & Raff, 2009). Nonetheless, the regulation of PCM dynamics remains incompletely understood.

Centrosomes are essential for early *Drosophila* embryogenesis, which proceeds through 14 rounds of rapid, synchronous nuclear cycles (NCs) prior to cellularization (Foe & Alberts, 1983). From NC 10–14, the embryo develops as a syncytial blastoderm, wherein thousands of nuclei and their associated centrosome pairs divide just under the embryonic cortex. Nuclear migration and divisions are coordinated by the centrosomes, and numerous mutations in centrosome-associated genes impair spindle morphogenesis, mitotic synchrony, genome stability, and embryonic viability (Deák et al., 1997; Freeman, Nüsslein-Volhard, & Glover, 1986; Sunkel & Glover, 1988). As in many organisms, the early development of the *Drosophila* embryo proceeds through a period of transcriptional quiescence and is supported by a maternal supply of mRNA and proteins (Vastenhouw, Cao, & Lipshitz, 2019). Thus, PCM dynamics apparent in early embryos rely upon post-transcriptional mechanisms.

Over a decade ago, a high-throughput screen for mRNAs with distinct subcellular locations in syncytial *Drosophila* embryos uncovered a subset of mRNAs localizing at or near spindle poles (Lécuyer et al., 2007). Many of the centrosome-enriched transcripts identified in that screen encode known centrosome regulators, including *cyclin B* (*cyc B*) and *pericentrin-like protein* (*plp*) (Dalby & Glover, 1992; Martinez-Campos, Basto, Baker, Kernan, & Raff, 2004; Raff, Whitfield, & Glover, 1990). These findings raise the possibility that RNA localization, translational control, and other post-transcriptional regulatory mechanisms contribute to centrosome activity and/or function. Consistent with this idea, RNA is known to associate with centrosomes in diverse cell types, including early embryos (*Drosophila, Xenopus*, zebrafish, and mollusk), surf clams, and cultured mammalian cells (Alliegro & Alliegro, 2008; Alliegro, Alliegro, & Palazzo, 2006; Blower, Feric, & Heald, 2007; Lambert & Nagy, 2002; Lécuyer et al., 2007; Raff et al., 1990; Sepulveda et al., 2018). The functional consequences and the mechanisms that regulate centrosome-localized RNA remain little understood, however (Marshall & Rosenbaum, 2000; Ryder & Lerit, 2018).

Here, we report that multiple RNA transcripts dynamically localize to centrosomes in *Drosophila* early embryos. We show these RNAs localize in unique patterns, with some RNAs forming higher-order granules, while others enrich around centrosomes as individual molecules. We further demonstrate that some RNAs enrich at centrosomal subdomains, such as the centrosome flares, which extend from interphase centrosomes and define the PCM scaffold (Lerit et al., 2015; Megraw, Kilaru, Turner, & Kaufman, 2002; Richens et al., 2015). We identify one centrosomal RNA, *centrocortin (cen)*, which forms micron-scale granules that localize asymmetrically to centrosomes. We further define the mechanisms underlying *cen* granule formation and function. We find that *cen* granules include Cen protein and the translational regulator fragile-X mental retardation protein (FMRP), the ortholog of the Fragile X Syndrome-related RNA-binding protein encoded by the *fmr1* gene. Our data show FMRP regulates both the localization and steady-state levels of *cen* RNA and protein. Moreover, we find that reducing *cen* dosage is sufficient to rescue the mitotic spindle defects associated with *fmr1* loss. Finally, we show that mislocalization of *cen* RNA prevents the localization of Cen protein to distal centrosomes and is associated with disrupted embryonic nuclear divisions.

## Results

### Quantitative analysis of mRNA distributions relative to *Drosophila* early embryonic centrosomes

A previous genome-wide screen identified a cohort of mRNAs showing apparent localization near spindle poles (Lécuyer et al., 2007). To quantitatively assess transcript localization to centrosomes, we combined single molecule fluorescence *in situ* hybridization (smFISH) with direct visualization of centrosomes. smFISH permits precise subcellular localization of individual RNA molecules, an important feature when determining enrichment at a relatively small target, such as the centrosome (Raj, van den Bogaard, Rifkin, van Oudenaarden, & Tyagi, 2008). For this analysis, we focused on syncytial embryos in NC 13, as their relatively prolonged interphase facilitates the collection of sufficient samples for quantification (Foe & Alberts, 1983). We used GFP-Centrosomin (GFP-Cnn) expressed under endogenous regulatory elements to label centrosomes (Lerit et al., 2015). Cnn is a core component of the centrosome scaffold required for the organization of the PCM that defines the outer edge of the centrosome (Conduit et al., 2010; Conduit et al., 2014; Megraw, Li, Kao, & Kaufman, 1999). Among the candidate RNAs reported to localize near spindle poles, we selected five for investigation based on prior data implicating their protein products in centrosome regulation and/or cell division: *cyc B, plp, small ovary (sov), partner of inscuteable (pins)*, and *cen* (Lécuyer et al., 2007; Fig. 1 Supplemental Table 1).

To examine patterns of RNA localization, we developed an automated custom image analysis pipeline that calculates the distribution of RNA transcripts relative to the distance from the centrosome (Fig. 1 Supplement 1A; see Methods). Briefly, smFISH signals and centrosomes were segmented, and the distances between individual RNA objects and the closest centrosome were measured (Fig. 1 Supplement 1B, C). This analysis allowed us to calculate the cumulative distribution of mRNA molecules relative to their distance from the surface of a centrosome (Fig. 1 Supplement 1D). We define mRNAs residing within 1 μm from the centrosome surface as pericentrosomal, or centrosome-enriched, because centrosomes extend dynamic Cnn-rich flares that rapidly sample this volume (Lerit et al., 2015; Megraw et al., 2002; Mennella et al., 2012). Among these localized mRNAs, those residing at 0 μm overlap with the centrosome (arrowheads, Fig. 1 Supplement 1C, D).

Several prior studies noted an enrichment of *cyc B* mRNA in the spindle pole region of syncytial *Drosophila* embryos (Dalby & Glover, 1992; Raff et al., 1990; Vardy & Orr-Weaver, 2007). Therefore, we initially investigated the localization of *cyc B* relative to a non-localizing control RNA, *gapdh*, to validate our quantitative imaging approach. Consistent with prior reports, we observed that *cyc B* was particularly abundant at the posterior pole (Raff et al., 1990). However, for the purposes of this study, all measurements were made in the somatic region at approximately 50% egg-length unless otherwise noted. To monitor cell cycle-dependent changes in RNA distribution, centrosome enrichments were calculated during interphase and metaphase. As expected, we found that *gapdh* was dispersed as single molecules throughout the cytoplasm (Fig. 1A, B), and few *gapdh* transcripts resided near centrosomes despite high levels of expression (Fig. 1C)(Graveley et al., 2011). By contrast, more *cyc B* transcripts localized in proximity to centrosomes, particularly during interphase (Fig. 1D, E). Approximately 2-fold more *cyc B* was enriched near centrosomes relative to *gapdh* (Fig. 1F and Fig 1. Supplement 2A, B).

**Figure 1.**
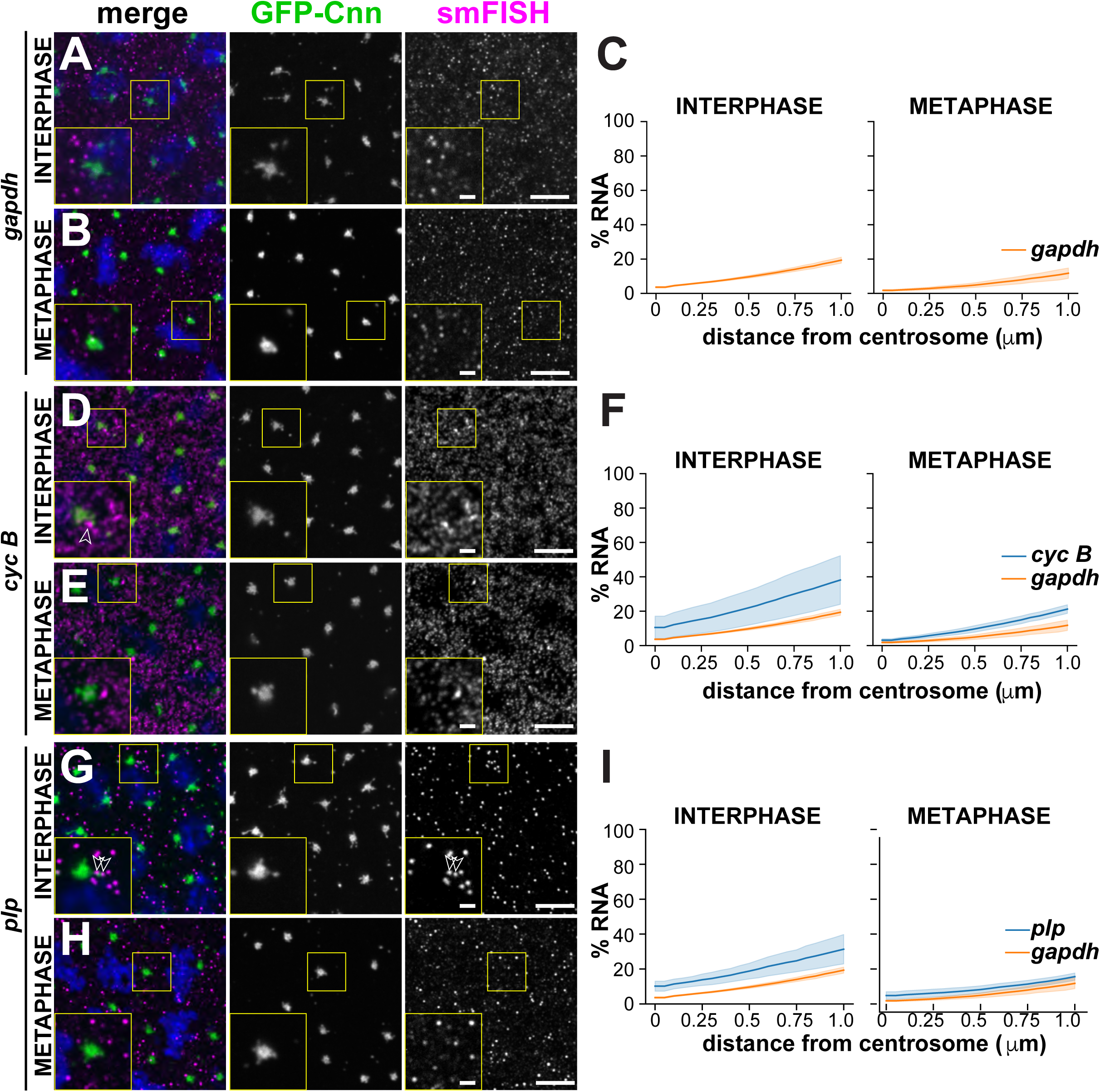
Quantitative localization of mRNA to centrosomes. Maximum intensity projections showing smFISH for the indicated RNAs (magenta) in interphase and metaphase NC 13 embryos expressing the centrosome marker GFP-Cnn (green). Nuclei are labeled with DAPI (blue). Boxed regions are enlarged in insets. Open arrowheads mark enrichments of *cyc B* and *plp* mRNAs near the PCM. Quantification of the cumulative percentage of RNA located within 1 μm from the centrosome surface is shown to the right and plotted as mean (dark line) ± S.D. (shading). (A–C) *gapdh*, (D–F) *cyc B*, and *plp* (G–I). See Fig. 1 Supplemental Table 2 for details regarding the number of embryos, centrosomes, and RNA objects quantified per condition. Scale bars: 5 μm and 1 μm (insets).

In interphase embryos, some *cyc B* smFISH signals appeared brighter and larger, suggesting that multiple *cyc B* transcripts clustered into higher order structures, hereafter referred to as RNA granules, near centrosomes (arrowhead, Fig. 1D). Quantification of the proportion of total RNA residing within granules, defined as an overlapping cluster of four or more mRNAs (Little, Sinsimer, Lee, Wieschaus, & Gavis, 2015), confirmed that more *cyc B* RNAs resided within pericentrosomal granules than *gapdh* (Fig. 1 Supplement 3A; Fig. 1 Supplemental Table 2). These findings demonstrate the utility of our analysis pipeline to quantitatively define RNA enrichments at centrosomes. Moreover, our data suggest that *cyc B* localization to centrosomes is regulated by granule formation and cell cycle progression.

### Multiple mRNAs are enriched at centrosomes in a cell-cycle dependent manner

We next investigated the localization of *plp* mRNA, as PLP protein cooperates with Cnn to mediate centrosome scaffolding (Lerit et al., 2015; Richens et al., 2015). Recently, orthologous *PCNT* transcripts were shown to be localized to centrosomes in zebrafish embryos and cultured mammalian cells, specifically during early mitosis (Sepulveda et al., 2018). In contrast, we found that *plp* transcripts frequently overlap with centrosomes during interphase (Fig. 1G–I; Fig 1. Supplement 2C). Specifically, *plp* was 1.6-fold enriched within 1 μm of centrosomes in interphase embryos relative to *gapdh*, yet only 1.3-fold enriched in metaphase embryos (Fig. 1I). We also noted that a subset of *plp* RNA (21.6% of total *plp* transcripts) localized to pericentrosomal granules in interphase (Fig. 1 Supplement 3B). By contrast, only 6.0% of *plp* transcripts in metaphase embryos were contained in pericentrosomal granules (Fig. 1 Supplement 3B; Fig. 1 Supplemental Table 2). These data reveal that *plp* mRNA enriches within granules at centrosomes specifically in interphase, coincident with the formation of centrosome flares containing PLP protein (Lerit et al., 2015), hinting that aspects of *plp* post-transcriptional regulation may be differentially regulated over the cell cycle.

We similarly analyzed the localization of *pins* and *sov* mRNAs relative to centrosomes. *pins* localized throughout the cytoplasm with only modest enrichments near centrosomes (Fig. 1 Supplement 2D and Fig. 1 Supplement 4A–C). Likewise, little *pins* is organized into RNA granules (Fig. 1 Supplement 3C; Fig. 1 Supplemental Table 2). In contrast, we found *sov* mRNA enriched at centrosomes (Fig. 1 Supplement 2E and Fig. 1 Supplement Fig. 4D–E), as previously noted (Lécuyer et al., 2007). smFISH highlights the propensity for *sov* mRNA to localize along centrosomal flares in interphase embryos (arrowheads, inset, Fig. 1 Supplement 4D). Consistent with these observations, over 20% of *sov* transcripts overlapped with interphase centrosomes (0 μm, Fig. 1 Supplement 4F), and ∼40% resided within 1 μm (1.9-fold enriched relative to *gapdh*; Fig. 1 Supplemental Table 2). Although centrosome-enrichment of *sov* is halved during mitosis (∼20% within 1 μm, Fig. 1 Supplement 4F), it was still 1.8-fold more enriched than *gapdh* (Fig. 1 Supplemental Table 2). Similarly, the proportion of *sov* within granules decreases upon mitotic onset (Fig. 1 Supplement 3D).

In sum, these findings reveal common and unique features of centrosome-localized mRNAs within *Drosophila* embryos. Generally speaking, localized mRNAs tend to be more enriched at centrosomes during interphase as compared to metaphase, although the magnitude of these changes varies by RNA. We also note increased biological variability in the proximity of mRNA to interphase centrosomes (e.g., compare error bars for interphase vs. mitosis, Fig 1F and I; Fig 1 Supplement Fig 2). These trends likely reflect the dynamic properties of the interphase centrosome, which extends protrusive flares to facilitate its expansion after mitotic exit (Lerit et al., 2015; Megraw et al., 2002). These data also suggest that RNA localization to centrosomes may be dynamic as well. RNA residence within granules is also more prevalent during interphase, similarly suggesting that aspects of RNA granule formation are cell cycle regulated.

### Dynamic regulation of micron-scale *cen* RNA granules

We next investigated the localization of *cen*, which was previously shown to be required for normal nuclear divisions in the *Drosophila* early embryo (Kao & Megraw, 2009). Unlike the other RNAs we investigated, the majority of *cen* was enriched at centrosomes (arrow, Fig. 2A). Throughout NC 13, we found that *cen* formed micron-scale granules, consistent with a recent report (Fig. 2A, B)(Bergalet et al., 2020). Demonstrating specificity, these signals were not detected in *cen* null mutant samples (Fig. 2 Supplement 1A). During interphase, these granules overlapped asymmetrically with a single centrosome (arrow, Fig. 2A). Further analysis revealed *cen* granules preferentially associate with mother centrosomes (Fig. 2 Supplement 1B, C). In metaphase, however, *cen* granules appeared less tightly associated with centrosomes (Fig. 2B). Quantification revealed that more than 50% of *cen* transcripts in NC 13 interphase embryos overlapped with centrosomes (resided at 0 μm), and over 80% of *cen* localized within 1 μm of a centrosome (Fig. 2C and Fig. 2 Supplement 2A). In metaphase, this enrichment is reduced (Fig. 2C). However, in both interphase and metaphase embryos, *cen* was approximately 4-fold more enriched at centrosomes relative to *gapdh* (Fig. 1 Supplemental Table 2). We further noted that fewer *cen* transcripts were detected in granules within 1 μm of a centrosome, dropping to 48% in metaphase from 75% in interphase (Fig. 2D; Fig. 2 Supplement 2B; and Fig. 1 Supplemental Table 2). These data demonstrate that *cen* forms micron-scale granules that localize to centrosomes in a cell cycle-dependent manner. These granules frequently overlap with centrosomes, resulting in a bulk enrichment of *cen* mRNA at centrosomes.

**Figure 2.**
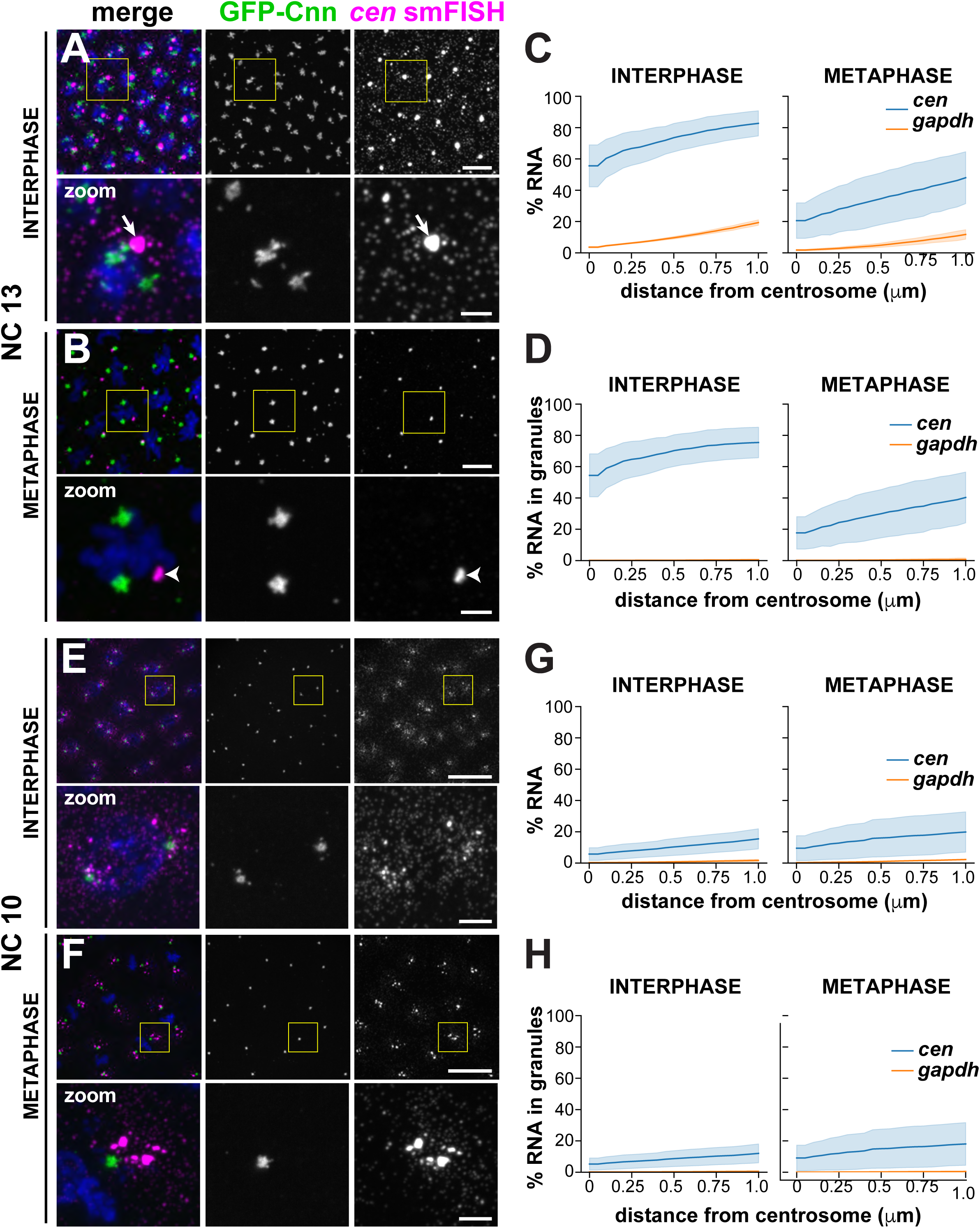
*cen* mRNA localizes to centrosomes in micron-scale granules that are cell cycle and developmentally regulated. Maximum intensity projections showing *cen* smFISH (magenta) in interphase and metaphase embryos expressing GFP-Cnn (green). Boxed regions are enlarged below (zoom). (A–D) Distribution of *cen* mRNA during NC 13. (A) During interphase, *cen* resides within a large granule asymmetrically localized to a single centrosome (arrow). (B) During metaphase, the *cen* granule is displaced from the centrosome (arrowheads). Quantifications show (C) the cumulative percentage of *cen* located within 1 μm from the centrosome surface and (D) the cumulative percentage of *cen* residing within RNA granules (≥ 4 overlapping RNAs) within 1 μm from the centrosome surface. During interphase, the majority of *cen* resides at the centrosome surface (0 μm) within RNA granules. (E–H) Distribution of *cen* mRNA during NC 10. (E) Interphase embryos show *cen* mRNA localized symmetrically and primarily as single molecules near centrosomes. (F) *cen* often resides within RNA granules in mitotic embryos. (G) Plot shows the cumulative distribution of *cen* located within 1 μm from the centrosome surface. (H) Plot shows the cumulative percentage of *cen* within RNA granules. Note the similarity to the cumulative distribution plot in (G), indicating that the majority of *cen* located within 1 μm of the centrosome is contained within granules. See Fig. 1 Supplemental Table 2 for details. Data are plotted as mean ± S.D. Scale bars: 10 μm and 2.5 μm (insets).

The strong enrichment of *cen* within pericentrosomal granules prompted us to investigate the developmental timing of their formation. In NC 10 embryos, the timepoint at which the syncytial nuclei first reach the cortex, we found that *cen* predominantly existed in single molecules radiating in a gradient from centrosomes (Fig. 2E). Entry into mitosis correlated with formation of larger *cen* granules that were closely apposed and symmetrically distributed to the two centrosomes (Fig. 2F). Similarly, the percentage of *cen* transcripts localized within 1 μm of a centrosome increased from ∼15% in interphase to nearly 20% in metaphase embryos (Fig. 2G; Fig. 2 Supplement 2C). Concordantly, the amount of *cen* RNA within pericentrosomal granules also increased, from 12% in interphase to 18% in metaphase (1 μm, Fig. 2H; Fig. 2 Supplement 2D; and Fig. 1 Supplemental Table 2). These data indicate that the formation of *cen* granules is entrained with the cell cycle and correlates with the initiation of cortical nuclear divisions. Our finding that *cen* RNA persists in a granular structure during interphase of NC 13 suggests the capacity for *cen* granule formation or maintenance is additionally regulated developmentally. Fewer granules are observed in younger embryos, which may be a feature of the abridged nature of those nuclear division cycles.

### The *cen* granule contains Cen protein, yet is dispensable for translation

To gain insight into the regulation and function of *cen* granules, we first investigated granule content. Recent work uncovered that *cen* granules contain Cen protein, and some *cen* granules represent sites of local translation (Bergalet et al., 2020). We similarly noted a strong coincidence of *cen* RNA and protein, confirming that Cen protein is abundant in *cen* granules (Fig. 3A, B).

**Figure 3.**
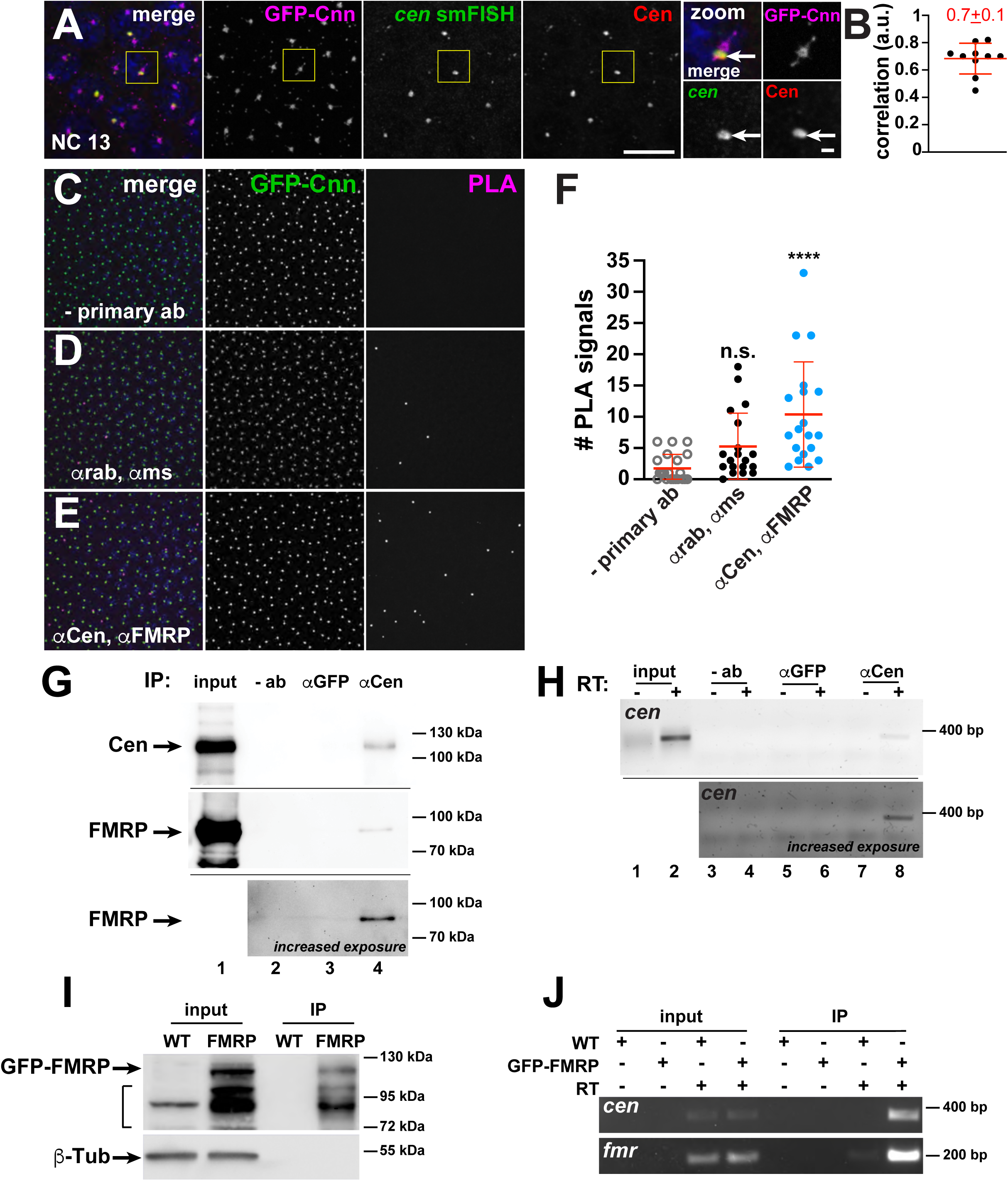
Composition of the *cen* granule. (A) Maximum intensity projection of a NC 13 embryo expressing GFP-Cnn (magenta) showing colocalization of *cen* mRNA (green) and protein (red). Boxed region is enlarged to the right (zoom); arrows highlight a *cen* granule. (B) Chart displays the Pearson’s correlation coefficient for colocalization between *cen* smFISH and anti-Cen signals (a.u., arbitrary units). Each dot represents a single measurement from N=10 NC 13 embryos; mean ± S.D. is shown (red). (C–E) Maximum intensity projections of NC 14 embryos expressing GFP-Cnn (green) with PLA signals (magenta) from the specified antibodies. (C) No primary antibodies, (D) control rabbit (rab) anti-Cnn and mouse (ms) anti-GFP antibodies, and (E) rabbit anti-Cen and mouse anti-FMRP antibodies. (F) Each dot shows the number of PLA signals counted within a single embryo within the field-of-view, ∼4,430 μm^2^, from N=21 embryos using no primary antibodies, N=19 embryos using control anti-rabbit and anti-mouse antibodies, and N=19 embryos using rabbit anti-Cen and mouse anti-FMRP antibodies; n.s. not significant; **** P ≤ 0.0001 by the Kruskal-Wallis test followed by Dunn’s test. (G) Immunoblot from anti-Cen immunoprecipitation using 1-3 hr embryonic extracts. Lane 1, 10% input; lane 2, no antibody (- ab)/empty beads; lane 3, control rabbit anti-GFP antibody; and lane 4, rabbit anti-Cen antibody. Cen pulls down itself (top) and FMRP (middle and bottom). The bottom blot shows an increased exposure to highlight the FMRP band; note, lane 1 was cropped due to over-saturated signal. (H) RNA-immunoprecipitation where RT-PCR reactions were run in the presence (+) or absence (-) of reverse transcriptase (RT). Lanes 1 and 2, 10% input; lanes 3 and 4, no antibody (- ab)/empty beads; lanes 5 and 6, control rabbit anti-GFP antibody; and lanes 7 and 8, rabbit anti-Cen antibody. The middle image shows an increased exposure to highlight the *cen* band; note, lanes 1 and 2 were cropped due to over-saturated signal. (I) Immunoblots from FMRP-GFP immunoprecipitation using 0-2 hr WT or FMRP-GFP embryonic extracts and GFP-Trap beads probed with rabbit anti-GFP (top) and mouse anti-β-Tub antibodies (bottom). GFP pulls out FMRP-GFP. Bracket denotes bands representing nonspecific and/or degradation products. (J) RNA-immunoprecipitation from GFP-Trap beads detects *cen* and the positive control, *fmr* (Ling, Fahrner, Greenough, & Gelfand, 2004). Scale bars: 10 μm and 1 μm (insets).

Previous work demonstrated Cen interacts directly with the centrosome scaffold protein, Cnn. In addition, a point mutation in Cnn, the *cnn*^*B4*^ allele, was sufficient to disrupt binding between Cnn and Cen and, consequently, Cen protein localization to centrosomes (Kao & Megraw, 2009). To test whether the centrosome scaffold is required for the localization of *cen* RNA to centrosomes, we examined if *cen* RNA localized to granules in *cnn*^*B4*^ mutants. We found that *cen* no longer formed granules in *cnn*^*B4*^ embryos and instead appeared dispersed throughout the cytoplasm as single molecules (Fig. 3 Supplement 1A). This behavior subsequently allowed us to test if *cen* granules were required for Cen translation. We observed no difference in the levels of Cen protein in 0-2 hour wild-type (WT) control versus *cnn*^*B4*^ mutant embryos, suggesting that the *cen* granule is not required for *cen* translation (Fig. 3 Supplement 1B, B’). These data support a model where the centrosome scaffold contributes to the formation of the *cen* granule, likely via associations between Cen and Cnn.

### FMRP associates with *cen* granules

RNA granules are diverse structures, and RNA-binding proteins are crucial for their formation and function (Singh, Pratt, Yeo, & Moore, 2015). Therefore, to provide mechanistic insight into the regulation of the *cen* granule, we assayed the centrosomal localization of a few candidate RNA-binding proteins, including Maternal expression at 31B (Me31B), Pumilio (Pum), Egalitarian (Egl), Orb2, and FMRP (Deshpande, Calhoun, & Schedl, 2006; Dienstbier, Boehl, Li, & Bullock, 2009; Gamberi, Johnstone, & Lasko, 2006); Fig. 3 Supplement 2A–E). Among these, a subset of FMRP puncta overlapped with centrosomes and *cen* granules (Fig. 3 Supplement 2E (arrowheads) and F (dashed circle)).

To further investigate the relationship between Cen and FMRP, we used a proximity ligation assay (PLA), which detects protein interactions when two primary antibodies bind antigens within a 40 nm threshold (Söderberg et al., 2006). In control experiments without Cen and FMRP antibodies, we rarely detected PLA signals in NC 14 embryos (Fig. 3C, D, and F). However, we detected a significant increase in PLA signals with Cen and FMRP antibodies (P<0.0001; Fig. 3E, F), indicating that subsets of these proteins reside in close physical proximity.

Finally, we biochemically probed *cen*-interacting factors. We isolated endogenous Cen protein complexes from early embryos by immunoprecipitation and found FMRP specifically associates with Cen (Fig. 3G). We similarly co-isolated *cen* RNA from Cen immunoprecipitates (Fig. 3H). Moreover, FMRP pulls down *cen* mRNA (Fig 3I, J). Taken together, we conclude that the *cen* granule represents a ribonucleoprotein (RNP) complex comprising several protein constituents, including Cen and FMRP. These data also hint that FMRP may mediate aspects of *cen* regulation.

### FMRP functions as a negative regulator of *cen* RNA granule formation and localization to centrosomes

FMRP is a multifunctional RNA-binding protein implicated in RNA localization, stability, and translational regulation (Banerjee, Ifrim, Valdez, Raj, & Bassell, 2018). To determine if FMRP contributes to *cen* regulation, we first compared the pericentrosomal localization of *cen* RNA and protein in control versus *fmr1* null mutant embryos expressing the PCM marker γ-Tubulin-GFP (γ-Tub-GFP). In control NC 10 interphase embryos, *cen* RNA was dispersed in predominantly single molecules near centrosomes, as we previously noted (Fig. 4 Supplement 1A). In *fmr1* embryos, however, *cen* RNA localized to granules of heterogenous size that clustered near centrosomes (Fig. 4 Supplement 1B), resulting in enhanced enrichment of *cen* near centrosomes (Fig. 4 Supplement 1E). In NC 10 metaphase embryos, *cen* formed small granules near centrosomes in control embryos (Fig. 4 Supplement 1C), but appeared to form larger granules in *fmr1* mutants (Fig. 4 Supplement 1D). Quantification revealed that 1.6-fold more *cen* was contained in granules localized within 1 μm of a centrosome in *fmr1* interphase embryos relative to controls (Fig. 4 Supplement 1F). In contrast, relatively similar levels of *cen* were contained in granules in *fmr1* embryos relative to controls during metaphase (Fig. 4 Supplement 1F; Fig. 4 Supplement 2A, B; Fig. 4 Supplemental Table 1). These data demonstrate that *cen* forms granules precociously in *fmr1* embryos, suggesting that FMRP normally limits *cen* localization to centrosomes.

We next investigated the contribution of FMRP to *cen* localization at later stages of development. In control NC 13 interphase embryos, *cen* formed micron-scale granules of heterogenous size (Fig. 4A). These pericentrosomal granules were larger in *fmr1* embryos (Fig. 4B), with 46% of *cen* transcripts overlapping with the centrosome in controls, compared to 68% in *fmr1* mutants (a 1.5-fold increase over WT; Fig. 4E; Fig. 4. Supplement 2C; Fig. 4 Supplemental Table 1). *fmr1* mutants also had more *cen* contained in granules during interphase, suggesting *cen* granule formation and/or localization was deregulated (Fig. 4F, Fig. 4 Supplement 2D). During metaphase, distributions of *cen* mRNA within *fmr1* mutants were more similar to WT, hinting that FMRP may contribute to the cell cycle dependent regulation of *cen* localization (Fig. 4C–F; Fig. 4. Supplement 2C and D; and Fig. 4 Supplemental Table 1).

**Figure 4.**
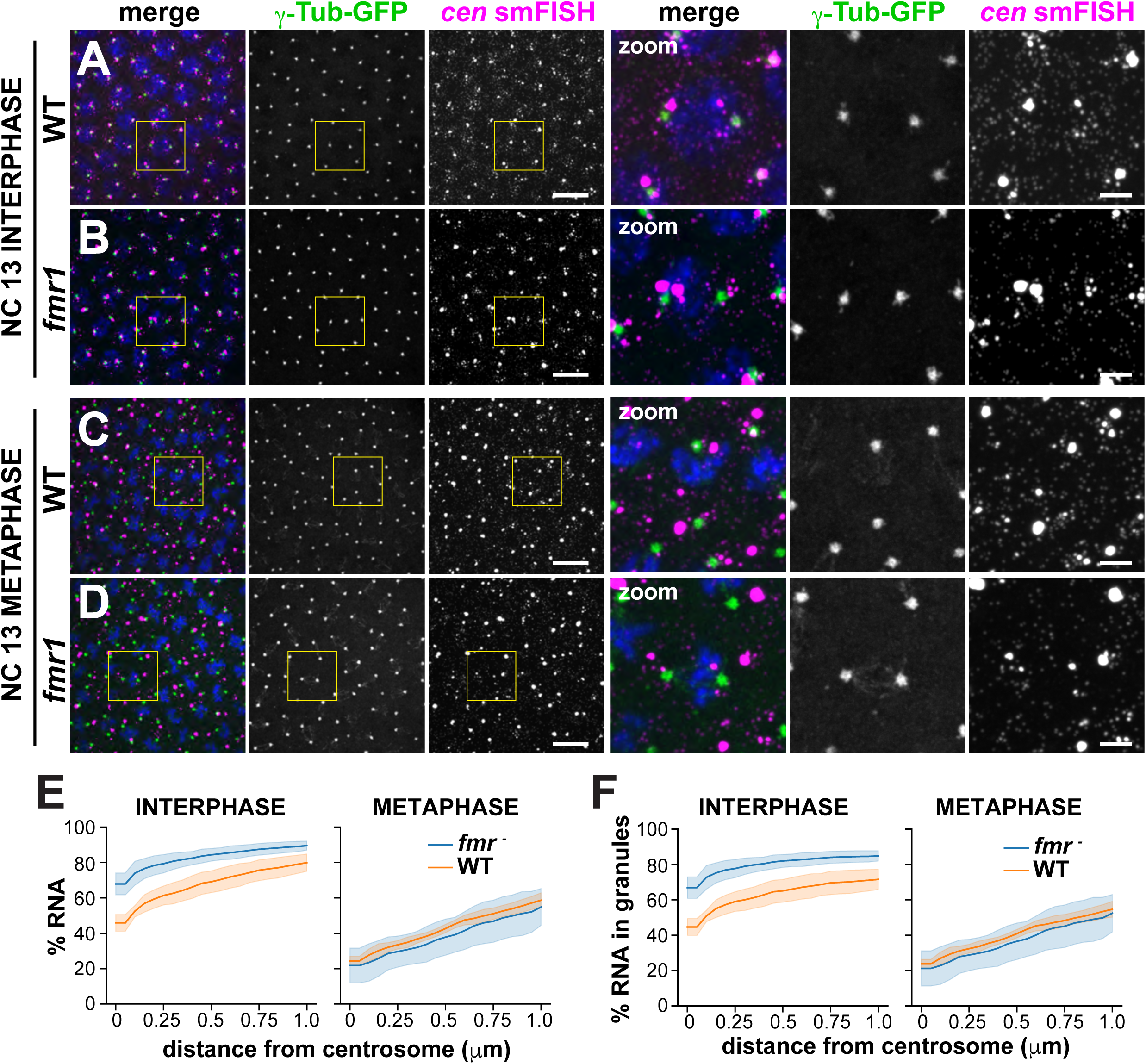
*Fmr1* regulates *cen* granule formation and size. Images show maximum intensity projections of WT or *fmr1* mutant NC 13 embryos expressing γTub-GFP and labeled with *cen* smFISH during (A and B) interphase or (C and D) mitosis. Boxed regions are enlarged to the right (zoom). (A) *cen* mRNA is typically packaged into a pericentrosomal granule in interphase control embryos. (B) *cen* granules are larger and less organized within *fmr1* embryos. (C) A control embryo showing *cen* granules displaced from mitotic centrosomes. (D) The distribution of *cen* mRNA within mitotic *fmr1* embryos resembles controls. (E) Graph shows the cumulative percentage of *cen* located within 1 μm of the centrosome surface in WT (orange) vs. *fmr1* mutant (blue) embryos. (F) Graph shows the cumulative percentage of *cen* found in RNA granules located within 1 μm of the centrosome surface. Data are plotted as mean ± S.D. See Fig. 4 Supplemental Table 1 for the number of embryos, centrosomes, and RNA objects quantified. Scale bars: 10 μm and 2.5 μm (insets).

These data show that loss of FMRP is associated with larger *cen* granules, which reside closer to and are more likely to overlap with centrosomes during interphase. We conclude that FMRP negatively regulates *cen* localization to centrosomes.

### FMRP regulates the abundance of *cen* RNA and protein

Since the early embryo is transcriptionally inactive for the first two hours of development (Anderson & Lengyel, 1979), the enhanced formation of *cen* granules in *fmr1* mutants could be attributed to changes in RNA localization, increased RNA stability, or both.

To test if FMRP contributes to *cen* RNA stability, we examined normalized *cen* RNA levels by qPCR. We found no significant change in *cen* RNA levels in *fmr1* vs. WT 0-1 hr embryos (P=0.07 by unpaired t-test; Fig. 5A). FMRP functions primarily as a translational repressor, and deregulation of FMRP targets in neurons is considered a significant driver of Fragile X Syndrome pathophysiology (Banerjee et al., 2018; Darnell, 2011). In 0–1 hr embryos, total levels of Cen protein were also unaffected by loss of *fmr1* (P=0.9 by unpaired t-test; Fig. 5B and B’). In contrast, within 1–3 hr embryos, *cen* RNA levels increased by 1.8–fold in *fmr1* mutants relative to controls (P<0.0001 by unpaired t-test; Fig. 5C). Similarly, 1–3 hr *fmr1* embryo extracts contained significantly more Cen protein than controls (3.7–fold increase in mutants relative to WT, P=0.03 by unpaired t-test; Fig. 5D and D’). Thus, while we found that both *cen* RNA and protein levels are increased in later stage *fmr1* embryos, the relative increase in Cen protein is nearly twice that observed for *cen* RNA. Taken together, these data suggest that FMRP contributes to *cen* RNA turnover and translational repression. The finding that younger *fmr1* mutant embryos show precocious and enhanced *cen* granule formation, despite WT levels of *cen* RNA, argues that changes in *cen* RNA localization and expression levels may be uncoupled and suggests that FMRP contributes to multiple aspects of *cen* RNA post-transcriptional regulation, either directly or indirectly.

**Figure 5.**
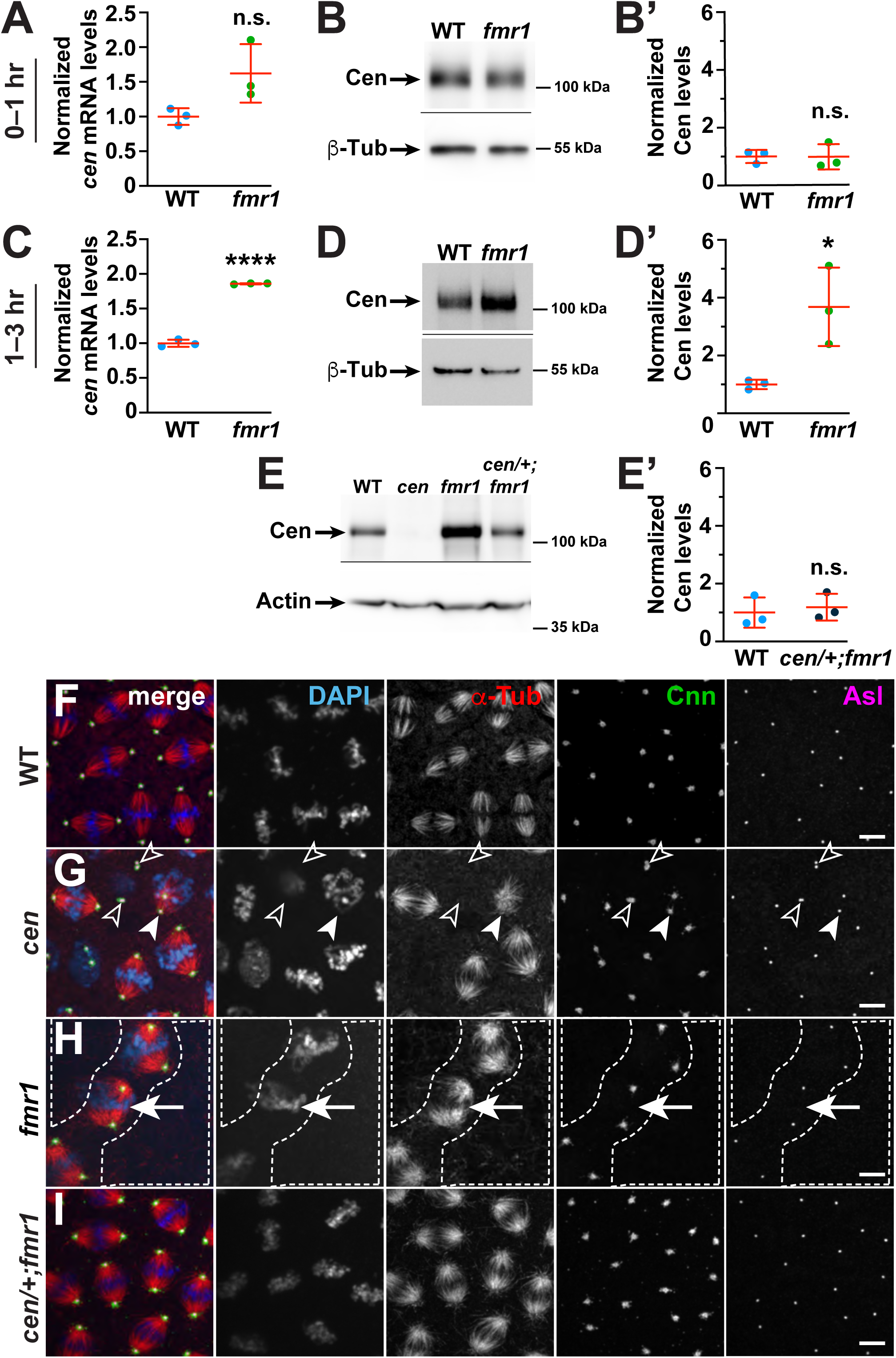
FMRP regulates *cen* to ensure error-free mitosis. (A) Levels of *cen* RNA were normalized to *RP49* as detected by qPCR from 0–1 hr embryonic lysates. (B) Immunoblots show Cen protein content relative to the β-Tub loading control from 0-1 hour embryonic extracts and are quantified in (B’). (C) Normalized levels of *cen* RNA from 1–3 hr embryos. (D) Immunoblots show Cen protein content relative to the β-Tub loading control in 1-3 hour embryonic extracts and are quantified in (D’). (E) Immunoblots show Cen protein content relative to actin loading control in 1-3 hour embryonic lysates from the indicated genotypes and are quantified in (E’). For (A–E’), data are normalized to the mean relative expression of the WT controls from N=3 biological replicates. (F–I) Maximum intensity projections of mitotic NC 11 embryos from the indicated genotypes showing immunofluorescence for α-Tub to label microtubules (red), Cnn labels PCM (green), and Asterless (Asl) labels centrioles (magenta). DAPI labels nuclei (blue). (F) WT embryos show normal, evenly spaced bipolar mitotic spindles. (G) Many *cen* embryos show spindle defects, including reduced microtubule organization and poorly condensed DNA (open arrowheads), as well as poorly separated centrosomes (closed arrowheads). (H) Spindle defects were common in *fmr1* mutants, as evidenced by massive nuclear fallout (dashed lines), as well as bent and disorganized spindles (arrows). (I) Hemizygosity for *cen* in the context of a *fmr1* background resulted in partial rescue of spindle defects and embryonic viability. n.s. not significant; *P< 0.05; **** P<0.0001 by unpaired t-test. Scale bars: 5 μm.

### *cen* and FMRP functionally interact to regulate cell division and embryonic viability

FMRP has established roles in progression through cell division. In neural progenitors, FMRP regulates proliferative capacity (Callan et al., 2010; Luo et al., 2010). In *Drosophila* embryos, loss of FMRP results in severe mitotic defects, including improper centrosome separation and loss of mitotic synchrony (Deshpande et al., 2006). In addition, many *fmr1* embryos form chromosome bridges or show evidence of lagging chromosomes or nuclear fallout, a developmental response to DNA damage resulting in the ejection of nuclei from the syncytial blastoderm cortex (Deshpande et al., 2006; Sullivan, Fogarty, & Theurkauf, 1993). Later in embryogenesis, loss of *fmr1* is also associated with defects in mitotic progression and cellularization (Monzo et al., 2006; Papoulas et al., 2010). Using hatch rate analysis as a measure of embryonic viability, we found that while *fmr1* mutants show an average of 6.3% unhatched embryos, *cen* hemizygosity partially restored viability (Table 1). Western blot analysis confirmed that *cen* hemizygosity normalized Cen protein levels in *fmr1* embryos (Fig. 5E and E’). These data are consistent with a genetic interaction between *cen* and *fmr1*; moreover, they implicate elevated Cen dosage as a driver of *fmr1*-mediated embryonic lethality.

To directly test if *cen* genetically modifies the mitotic defects observed in *fmr1* mutant embryos, we tabulated the incidence of abnormal microtubule spindles. Occasionally, even WT embryos contained aberrant microtubule spindles (3.7%, N=1/27 embryos; Fig. 5F). However, *cen* mutant embryos showed an increased rate of spindle errors (40.9%, N=9/22 embryos; Fig. 5G, arrowheads), consistent with prior observations (Bergalet et al., 2020; Kao & Megraw, 2009). Similarly, loss of *fmr1* was associated with high rates of spindle defects (76.1%, N=16/21 embryos; Fig. 5H, arrow). We also noted areas of lower nuclear density in *fmr* embryos, consistent with nuclear fallout (Fig. 5H, dashed lines). In contrast, reducing *cen* dosage in the context of the *fmr1* null background partially rescued the incidence of mitotic spindle defects (48.1%, N= 13/27 embryos; Fig 5I). Together, these data demonstrate that normal dosage of Cen is required for error-free mitosis and that the upregulation of *cen* in *fmr1-*null embryos contributes to an increased rate of spindle errors and embryonic lethality.

### Ectopic *cen* localization disrupts nuclear divisions

Our data support a model whereby the local concentration of *cen* contributes to proper cell cycle progression. To directly test this model, we engineered a chimeric RNA comprising the *cen* coding sequence and the *bicoid* (*bcd*) 3’UTR, previously shown to be sufficient to mislocalize target RNAs to the anterior pole (Macdonald & Struhl, 1988). For these experiments, we examined embryos from mothers expressing the *cen-bcd 3’UTR* transgene in the context of the *cen* null background (hereafter, *cen-bcd 3’UTR* embryos).

We first confirmed our transgenic construct successfully mistargeted *cen* RNA to the anterior. Pre-blastoderm *cen-bcd 3’UTR* embryos (aged ∼0–30 min), showed a crescent of *cen* RNA at the anterior pole (Fig. 6A). Immunofluorescence revealed that Cen protein is translated and also localized to the anterior of young *cen-bcd 3’UTR* embryos (Fig. 6A). Immunoblotting showed Cen protein is expressed in *cen-bcd 3’UTR* embryos at levels comparable to *fmr1* mutants (mean 3.3-fold increased relative to WT controls, P=0.03 by unpaired t-test; Fig. 6 Supplement 1A, A’). Given the restricted localization of *cen* to the anterior pole, the local concentration of *cen* mRNA and protein is expected to be significantly higher than normal.

**Figure 6.**
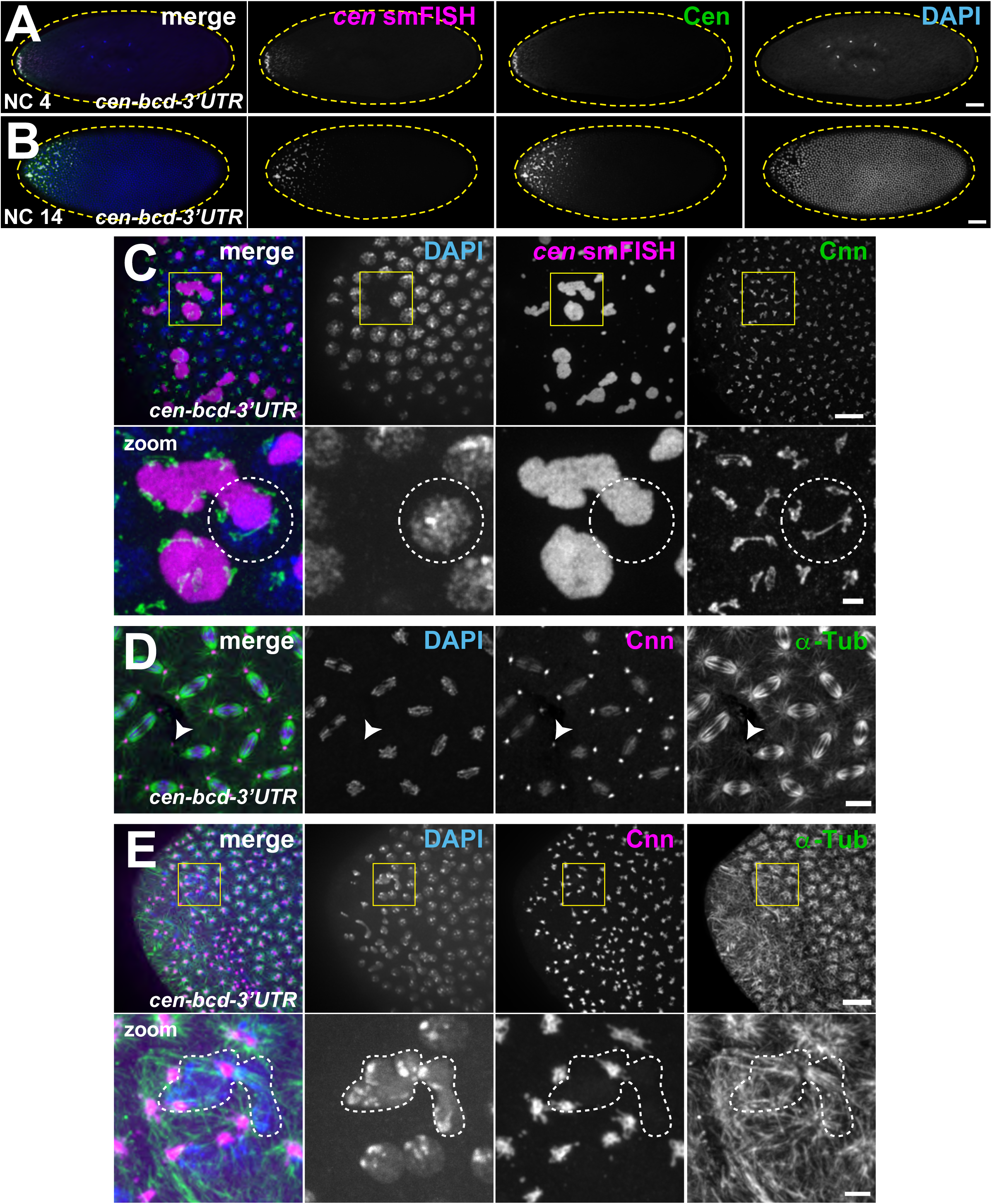
Ectopic localization of *cen* RNA disrupts nuclear divisions. Images show maximum intensity projections of *cen-bcd-3’UTR* embryos, which are progeny from females expressing the *pUASp*-*cen-bcd-3’UTR* transgene under the *maternal α-Tub GAL4* driver in the *cen* null background. (A and B) Low magnification images showing anterior localization of *cen* smFISH signals (magenta) costained with DAPI (blue) and anti-Cen antibodies (green). (A) NC 4 embryo showing a gradient of *cen* RNA and protein focused at the anterior pole. (B) NC 14 embryo showing the disruption of nuclear spacing at the anterior pole. (C) Ectopic localization of *cen* RNA to the anterior pole results in the formation of massive *cen* RNA granules (magenta) decorated by numerous centrosomes (Cnn, green). Boxed region is enlarged below (zoom); dashed circle highlights a nucleus and part of a *cen* RNP associated with supernumerary centrosomes. Nuclear fallout is evident by holes in the nuclear monolayer. (D) NC 12 embryo showing a mitotic spindle defect at ∼50% egg-length; arrowhead marks a detached centrosome. (E) NC 12 embryo showing extensive disruptions to microtubule organization (α-Tub, green) and centrosome positioning (Cnn, magenta) at the anterior pole. DAPI-labeled nuclei (blue) are often enlarged or dysmorphic (dashed lines). Clusters of anucleated centrosomes indicate nuclear fallout. Boxed regions show insets enlarged below (zoom). Scale bars: (A and B) 50 μm; (C–E) 10 μm and 2 μm (insets).

At the anterior of *cen*-*bcd-*3’UTR embryos, *cen* RNA and protein coalesced into RNPs, which were much larger than the typical *cen* granules observed in WT (Fig. 6B). These large RNPs were also prominent during NC 10, when *cen* normally exists as single molecules (Fig. 6 Supplement 1B). Through the use of reporter constructs, it was recently demonstrated that the *cen* coding sequence is sufficient for centrosome targeting (Bergalet et al., 2020). Consistent with this idea, the enlarged *cen* RNPs observed in *cen*-*bcd-*3’UTR embryos retained the ability to associate with centrosomes (dashed circle, Fig. 6C and Fig. 6 Supplement 1B). These data suggest that the normal temporal and spatial pattern of *cen* RNA localization requires sequence or structural elements encoded within the native *cen* 3’UTR.

We did not observe *cen* RNA or protein localized to distal centrosomes in *cen-bcd 3’UTR* embryos, suggesting that localization elements within the *bcd* 3’UTR confine *cen* localization to the anterior pole. Moreover, this finding suggests that proper localization of *cen* mRNA is required for Cen localization to centrosomes. The restricted localization of *cen* mRNA and protein to the anterior pole within *cen-bcd 3’UTR* embryos allowed us to test whether *cen* was required locally for error-free mitosis. Examination of mitotic spindles at ∼50% egg-length within *cen-bcd 3’UTR* embryos revealed an increased rate of microtubule spindle defects (47.6%, N=10/21 embryos; Fig. 6D), indicating that *cen* functions locally to support normal spindle morphogenesis.

Notably, the anterior pole of *cen-bcd 3’UTR* embryos showed lower nuclear density (i.e., nuclear fallout), dysmorphic nuclei, and mitotic asynchrony, showcasing significant disruption to nuclear divisions (Fig. 6B and E). To further characterize the underlying mechanisms responsible for the nuclear division defects observed at the anterior region of *cen*-*bcd-3’UTR* embryos, we examined their mitotic spindles. We found severe disruptions to microtubule organization in these embryos, as well as nuclei associated with supernumerary centrosomes (Fig. 6E). Quantification revealed that 85% of *cen*-*bcd-3’UTR* embryos showed spindle defects at the anterior (N=17/20 embryos). In contrast, spindle defects occurred less frequently in control embryos (N=2/21 embryos; similar results observed in N=3 independent replicates for both genotypes).

Given these phenotypes, we next examined embryonic viability. While *cen* mutant embryos show an elevated rate of unhatched embryos relative to controls, consistent with prior work (mean 10.7% unhatched; (Kao & Megraw, 2009), *cen-bcd 3’UTR* embryos showed increased lethality (mean 19.2%; P=0.049 relative to *cen* by unpaired t-test; Table 1). We propose a model in which the deregulated balance of Cen levels impairs mitotic spindle organization (Fig. 7). Collectively, our data suggest that temporal and spatial regulation of *cen* RNA at centrosomes is required for error-free mitosis and embryonic viability.

**Figure 7.**
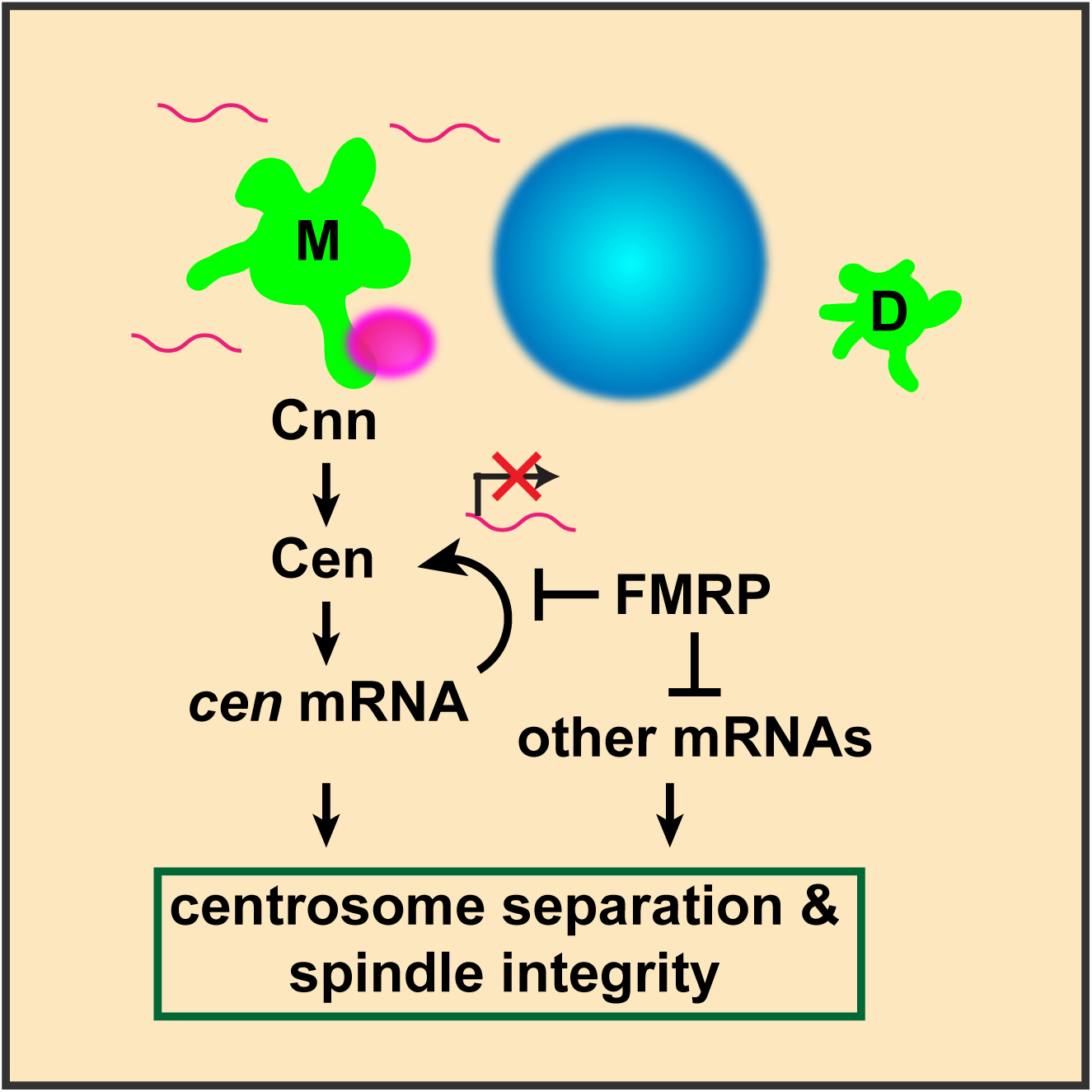
Model of FMRP-mediated *cen* mRNA localization and translational control at centrosomes. Diagram illustrates *cen* mRNA (magenta) recruitment to interphase centrosomes (green); nucleus is blue. A direct interaction between Cnn and Cen recruits Cen to the centrosome (Kao & Megraw, 2009). Cen protein is sufficient to recruit *cen* mRNA, and local translation of Cen creates a positive feedback loop, resulting in a concentrated, pericentrosomal enrichment of *cen* (Bergalet et al., 2020). We show that *cen* mRNA and protein form an immunoprecipitable complex, and they colocalize within micron-scale granules. We further show that the localization of *cen* mRNA to centrosomes, its organization into granules, the stability of *cen* mRNA, and its translation are all regulated by FMRP. Finally, our genetic epistasis work demonstrates that *cen* is an important target of FMRP required for centrosome separation, spindle morphogenesis, and error-free mitosis.

## Discussion

Centrosome-localized RNA has been described in a variety of organismal contexts, and while the conserved feature of mRNA at centrosomes hints at a biological function, the underlying physiological significance has remained unclear (Marshall & Rosenbaum, 2000; Ryder & Lerit, 2018). To begin to resolve this question, we systematically examined five transcripts predicted to enrich near spindle poles, and we quantitatively characterized their common and unique localization patterns in interphase and mitotic *Drosophila* embryos. We identified subsets of mRNAs showing centrosome enrichment in a cell cycle and developmentally regulated manner. These non-random variances in RNA distributions over time further imply biological relevance. We directly tested if RNA localization contributes to normal centrosome functions through in-depth studies with the model transcript *cen*. We identified FMRP as an RNA-binding protein required for the regulation of *cen* RNA localization, organization, and translational control. Further, we showed that reducing *cen* dosage ameliorates *fmr1*-dependent mitotic errors and embryonic lethality. We also directly tested the consequences of mistargeting *cen* mRNA. Mislocalization of *cen* mRNA to the anterior abrogated the normal localization of Cen to more distal centrosomes and disrupted spindle organization. Anterior mitotic divisions were also severely disrupted due to the increased local concentration of Cen. These studies suggest that a normalized local concentration of *cen* is essential for normal cell division and genome stability.

### Centrosomes as platforms for translational regulation

FMRP is a multifunctional RNA-binding protein with roles in translational repression, activation, RNA localization, and RNA stability (Darnell, 2011; Estes, O’Shea, Clasen, & Zarnescu, 2008; Greenblatt & Spradling, 2018; Pilaz, Lennox, Rouanet, & Silver, 2016). In humans, mutations in the gene encoding FMRP, *FMR1*, are the leading cause of heritable intellectual disability and autism. As a result, numerous high-throughput studies have identified putative RNA substrates, although surprisingly few of these have been validated (Santoro, Bray, & Warren, 2012). Our studies demonstrate that *cen* is regulated by FMRP, either directly or indirectly, and that titrating *cen* dosage is sufficient to partially restore embryonic viability in *fmr1* mutants. Consistent with direct regulation of *cen* by FMRP, the *cen* coding sequence contains six putative binding motifs for FMRP, according to RBPmap, an RNA-binding motif predictor (Paz, Kosti, Ares, Cline, & Mandel-Gutfreund, 2014). Moreover, human orthologs of *cen, CDR2* and *CDR2L*, were identified as direct FMRP targets by PAR-CLIP (photoactivatable ribonucleoside-enhanced crosslinking and immunoprecipitation) (Ascano et al., 2012). Deregulation of *CDR2* and *CDR2L* is associated with paraneoplastic cerebellar degeneration, indicating that their altered levels or activities contribute to neural degeneration (Albert et al., 1998; Corradi, Yang, Darnell, Dalmau, & Darnell, 1997). Our studies suggest *Drosophila cen* may serve as a valuable model to uncover mechanisms underlying FMRP-mediated regulation of *CDR2* and *CDR2L*.

The enhanced recruitment of *cen* to heterogeneously sized pericentrosomal granules, coupled with the increased production of Cen protein within *fmr1* mutants, led us to speculate that *cen* granules may be sites of local translation, as was recently proposed (Bergalet et al., 2020). However, disruption of *cen* granule formation, as in *cnn*^*B4*^ mutants, does not impair total Cen protein levels. This finding raises the possibility that Cen may be translated at alternate sites or that maternal stores of Cen obscure changes resulting from *cen* granule loss. Nonetheless, our data suggest that centrosomes serve as platforms for translation control, which may be positive or negative depending on the specific transcript and/or cell cycle stage. We propose a model, wherein *cen* granules are sites of Cen translational regulation (Fig. 7). Our data suggest that FMRP functions as a negative regulator of *cen*, limiting Cen expression and, consequently, *cen* granule size and bulk enrichment at centrosomes. In the absence of FMRP, Cen expression becomes deregulated and may help recruit additional *cen* mRNA molecules from the cytoplasm to enlarged pericentrosomal granules. An imbalance of Cen levels at centrosomes – either too little (as in *cen* mutants) or too much (as in *fmr* mutants or *cen*-*bcd-3’UTR* embryos) – impairs centrosome function/spindle integrity and embryonic viability. Given our finding that loss of *fmr* does little to *cen* localization in mitotic embryos, we speculate that FMRP normally represses *cen* during interphase. Cen expression may normally be derepressed upon mitotic onset to permit local translation.

### Differential enrichment of mRNAs on interphase centrosomes

A common trend emerging from our comparative analyses is the greater enrichment of RNA at centrosomes during interphase versus metaphase, as exemplified by *cen, cyc B, plp, and sov*. One possible explanation is the differential size of interphase centrosomes, which are significantly larger in *Drosophila* embryos due to the elaboration of extended centrosome flares, part of the architecture of the centrosome scaffold (Lerit et al., 2015; Megraw et al., 2002; Richens et al., 2015). This pattern contrasts with mammalian centrosomes, which are smaller in interphase and larger in mitosis (Lawo, Hasegan, Gupta, & Pelletier, 2012). According to this size model, a larger centrosome might nonspecifically recruit or dock additional RNAs simply due to the increased volume it occupies in the cell. We discount this model based on our finding that a highly expressed control transcript, *gapdh*, does not enrich at interphase centrosomes. This result also argues against the idea that centrosomes non-specifically recruit RNA molecules spuriously. Relatively few RNAs localize to centrosomes, while many others do not (Lécuyer et al., 2007; Raff et al., 1990). Here, we show the localization of centrosome-associated RNA is regulated in space and time.

Why do RNAs localize to interphase centrosomes? Recent work in mammalian cells proposed that some lengthy transcripts may be cotranslationally transported to centrosomes (Sepulveda et al., 2018). This model would account for contemporaneous recruitment and colocalization of centrosome RNA and proteins. If some centrosome transcripts utilize cotranslational transport, *sov* may prove to be an exception. Of the RNAs overlapping with the centrosome surface, *sov* was unique in that it appeared to preferentially dock along centrosome flares, localizing to the outer PCM zone. However, we do not detect Sov at centrosomes. Instead, Sov resides in the nucleus during interphase and is undetectable after nuclear envelope breakdown (Benner et al., 2019). These findings suggest that Sov is rapidly translocated into the nucleus. Live imaging of RNA transport and nascent protein synthesis is required to rigorously test the dynamics of RNA localization and local translation.

Another model that may account for enrichment of centrosome RNAs at interphase centrosomes is the possibility that RNA contributes to centrosome structure, perhaps as a component of the PCM scaffold itself. Recent work has suggested that phase transitions may contribute to PCM structure and function (Woodruff et al., 2017; Woodruff et al., 2015; Zwicker, Decker, Jaensch, Hyman, & Jülicher, 2014). A common principle of phase transitions is the association of intrinsically disordered proteins with specific RNA molecules to form non-membrane bound organelles with unique biophysical properties (Berry, Brangwynne, & Haataja, 2018). Centrosome-associated RNA may function as a physiological crowding agent contributing to phase transitions of the PCM. A related intriguing question raised by our work is if *cen* granules represent phase-separated domains. Cen protein contains multiple predicted intrinsically disordered domains, which is congruous with phase-separation (Ishida & Kinoshita, 2007). While we cannot rule out the contribution of all centrosome-enriched RNAs, our studies do not support a model for *cen* RNA contributing to centrosome structure. Mistargeting *cen* to the anterior cortex did not appear to disrupt the organization of distal centrosomes, for example.

Critically, disrupting the PCM scaffold is sufficient to inhibit *cen* granule formation. We previously showed that the PCM scaffold becomes progressively more structured during the prolonged interphases of later NCs (Lerit et al., 2015). Additionally, the mother centrosome organizes a larger PCM scaffold due to inherently greater levels of Cnn and PLP (Conduit et al., 2010; Lerit et al., 2015). Collectively, these features may account for the asymmetric localization of *cen* granules to mother centrosomes in late-stage syncytial embryos. These data lead us to conclude that the PCM scaffold organized by Cnn and PLP is upstream of the recruitment and organization of *cen* RNA granules (Fig. 7).

### Towards an understanding of the post-transcriptional regulation of centrosomal RNAs

Our finding that some population of most pericentrosomal RNAs organize into higher-order granules hints that these structures might represent regulatory RNPs. Many types of RNP granules form within cells, including stress granules, germ granules, P-bodies, etc., which all have unique functions and modes of assembly. The spatial proximity of multiple RNA molecules may facilitate intermolecular RNA interactions subsequently recognized by RNA-binding proteins (Van Treeck & Parker, 2018). While the FMRP-containing *cen* granule represents one such RNP, an area of active investigation in our lab is the functional characterization of other centrosomal RNAs. As the early *Drosophila* embryo is transcriptionally quiescent, post-transcriptional regulatory mechanisms, and especially translational control, are fundamentally important for proper centrosome regulation and function.

## Methods

### Fly stocks

The following *Drosophila* strains and transgenic lines were used: *y*^*1*^*w*^*1118*^ (Bloomington *Drosophila* Stock Center, BDSC #1495) was used as the WT control unless otherwise noted; *P*_*BAC*_*-GFP-Cnn*, which expresses Cnn tagged at the N-terminus with EGFP under endogenous regulatory elements (Lerit et al., 2015); *Ubi-GFP-γ-Tub23C* expresses *GFP-γ-Tub* under the *Ubiquitin* promotor (Lerit & Rusan, 2013); null *cen* mutant embryos derive from homozygous *cen*^f04787^ animals (BDSC #18805) (Kao & Megraw, 2009); null *fmr1* mutant embryos derive from *fmr1*^*Δ113M*^*/fmr*^*3*^ trans-heterozygotes (*fmr1*^*Δ113M*^ BDSC #67403) (Zhang et al., 2001); *fmr*^*3*^ gift from T. Jongens, UPenn) (Dockendorff et al., 2002); and hypomorphic *cnn*^*B4*^ mutants were a gift from T. Megraw (Florida State University). The *maternal α-Tub* promoter was used to control *GAL4* expression (*matGAL4*; BDSC #7063) to drive expression of *pUASp-cen-bcd-3’UTR* (this study). *FMRP-GFP* is a recombineered line expressing FMRP tagged at the C-terminus with GFP under endogenous regulatory elements (gift from M. Ramaswami, Trinity College Dublin) (Sudhakaran et al., 2014). In all experiments, mutant embryos represent progeny derived from mutant mothers to examine maternal effects. Flies were raised on molasses-based *Drosophila* medium, and crosses were maintained at 25°C in a light and temperature-controlled chamber.

### Construction of transgenic animals

To generate *pUASp-cen-bcd-3’UTR*, the *cen* coding sequence was PCR amplified using Phusion high-fidelity DNA polymerase from the cDNA clone LD41224 (*Drosophila* Genomics Resource Center (DGRC)) using the primers 5’-GCAGGCTCCGCGGCCGCCCCCTTCACCAG-G<u>ATG</u>GAGGAATCCAATCACGGTTC-3’ and 5’-GAAACTCTCTAACAGCCTCTCATCCAGG<u>T-TA</u>CTTTTGACGAAACTGATGATGATGACTC-3’. The *bcd-3’UTR* was PCR amplified using Q5 high-fidelity polymerase (New England Biolabs, M0491S) from genomic DNA using the primers 5’-GAGTCATCA-TCATCAGTTTCGTCAAAAG<u>TAA</u>CCTGGATGAGAGGCGTGTTAGAG-3’ and 5’-CTGGGTCG-GCGCGCCCACCCTTGTCTAGGTAGTTAGTCACAATTTACCCGAGTAGAGTAG-3’. The *cen* start and stop codons are underlined. The *cen-bcd-3’UTR* fusion was assembled and directionally cloned into the pENTR-D vector (Invitrogen) by Gibson assembly using 5-fold molar excess of the *bcd-3’UTR*. Sequence-verified single colony clones were shuttled into the destination vector pPW (UASp promoter) using the Gateway cloning system (Invitrogen). Transgenic animals were generated by BestGene, Inc.

### Embryonic hatch rate analysis

24-hr collections of eggs were collected on yeasted grape juice agar plates, transferred to fresh grape juice agar plates, and aged for 48-hr at 25 °C. Unhatched embryos were counted from a total of ∼600 embryos, and data presented are mean ± S.D. from 3 biological replicates.

### Immunofluorescence

Embryos were prepared for immunofluorescence as described in Lerit et al. 2015. Briefly, samples were fixed in paraformaldehyde, blocked extensively in BBT (PBS supplemented with 0.1% Tween-20 and 0.1% BSA, or 0.5% BSA for Asl staining), then incubated overnight at 4°C with primary antibodies diluted in BBT. The next day, samples were further blocked in BBT supplemented with 2% normal goat serum (NGS) and then incubated with secondary antibodies and DAPI for 2 hr at room temperature prior to mounting in AquaPoly/Mount mounting medium (VWR, 87001-902).

The following primary antibodies were used: rabbit anti-Cen (1:500; gift from T. Megraw, Florida State University) (Kao & Megraw, 2009), rabbit anti-Cnn (1:3500; gift from T. Megraw), guinea pig anti-Asl (1:3000; gift from G. Rogers, University of Arizona), mouse anti-*α*-Tub DM1*α* (1:500; Sigma, T6199), rabbit anti-Egl (1:2000; gift from R. Lehmann, New York University), rabbit anti-Pum (1:1000; gift from Martine Simonelig, Institute of Human Genetics, University of Montpellier), mouse anti-Orb2 (1:1000; Developmental Studies Hybridoma Bank (DSHB) clone 4G8), mouse anti-Me31b (1:3000; gift from A. Nakamura, Kumamoto University); and mouse anti-FMRP (1:10; DSHB clone 5A11).

Secondary antibodies and stains: Alexa fluor 488, 568, or 647 (1:500, Molecular Probes). DAPI was used at 10 ng/mL (Thermo Fisher).

### Detection of RNA by smFISH

smFISH experiments were adapted from manufacturer’s recommended protocols. All steps were performed with RNase-free solutions. Briefly, fixed and rehydrated embryos were washed in PBST (PBS plus 0.1% Tween-20) then washed in wash buffer (WB; 10% formamide and 2x saline sodium citrate (SSC) supplemented fresh each experiment with 0.1% Tween-20 and 2 μg/mL nuclease-free BSA (VWR, 0332-25G)). Embryos were then incubated with 100 μL of hybridization buffer (HB; 100 mg/mL dextran sulfate and 10% formamide in 2x SSC supplemented fresh each experiment with 0.1% Tween-20, 2 μg/mL nuclease-free BSA, and 10 mM ribonucleoside vanadyl complex (RVC; New England Biolabs, S1402S) for 10-20 minutes in a 37°C water bath. Stellaris smFISH probes conjugated to Quasar 570 dye (LGC Biosearch Technologies) were designed against the coding region for each gene of interest using the Stellaris RNA FISH probe designer and stored at −20 °C as stock solutions of 25 μM in nuclease-free water. See Fig. 1 Supplemental Table 3 for detailed information regarding probes. After pre-incubation in HB, embryos were incubated in a 37 °C water bath overnight in 25 μL of HB containing a 1:50 dilution of smFISH probe. The next morning, embryos were washed three times for 30 min each in pre-warmed WB, stained with DAPI for 1 hr at room temperature, washed with PBST, then mounted with Vectashield mounting medium (Vector Laboratories, H-1000). Slides were stored at 4 °C and imaged within 1 week.

For experiments where immunofluorescence was combined with smFISH, we adapted a protocol from Xu et al., 2015. Following an overnight incubation with smFISH probes, embryos were washed well in WB, followed by two 10 min washes in 2x SSC-0.1% Tween-20, and then four 10 min washes in PBST. Embryos were then blocked for two hours in blocking solution (PBS supplemented with 1 mg/mL nuclease-free BSA, 0.1% Tween-20, and 2 mM RVC, prepared fresh), then incubated overnight in primary antibodies at 4 °C. The next day, embryos were washed well in blocking solution, incubated with secondary antibodies and DAPI at room temperature, then washed in PBST prior to mounting in Vectashield.

### Proximity Ligation Assays

Proximity ligation assays were performed using the Sigma Duolink PLA kit (DUO92101) following manufacturer’s recommendations with minor modification. Fixed and rehydrated embryos were incubated in 1 drop (∼ 40 μL) of the Duolink blocking solution at 37 °C for 60 min without nutation. Embryos were then incubated in primary antibodies diluted in BBT (PBS, 0.1% Tween-20, and 0.5% BSA) overnight at 4 °C. The next day, embryos were washed with Duolink Wash Buffer A twice for 5 min, incubated with 40 μL of Duolink PLA probes diluted 1:5 in Duolink antibody diluent for 60 min at 37 °C, re-washed in Wash Buffer A, incubated with Duolink Ligase diluted 1:40 in 1x Duolink Ligation Buffer at 37 °C for 30 min, and then re-washed with Wash Buffer A. The amplification step was then performed using 0.5 μL of polymerase diluted in 40 μL of 1x amplification buffer at 37 °C for 100 minutes. Finally, embryos were incubated with DAPI diluted in Wash Buffer B for 15 min, washed twice in Wash Buffer B, and mounted in Vectashield. Slides were stored at −20 °C and imaged within 48 hours.

The following primary antibody pairs were used: rabbit anti-Cen (1:500) and mouse anti-FMRP (1:10, DSHB); negative controls included rabbit anti-Cnn (1:3500) and mouse anti-GFP (1:1000; DSHB clone 4C9) and no primary antibodies.

### Microscopy

Images were acquired on a Nikon Ti-E system fitted with a Yokogawa CSU-X1 spinning disk head, Hamamatsu Orca Flash 4.0 v2 digital CMOS camera, Perfect Focus system, and a Nikon LU-N4 solid state laser launch (15 mW 405, 488, 561, and 647 nm) using the following objectives: 100x 1.49 NA Apo TIRF oil immersion, 40x 1.3 NA Plan Fluor oil immersion, and 20×0.75 NA Plan Apo. This microscope was powered through Nikon Elements AR software on a 64-bit HP Z440 workstation.

### Image analysis

Images were assembled using Fiji (NIH) (Schindelin et al., 2012), Adobe Photoshop, and Adobe Illustrator software to separate or merge channels, crop regions of interest, generate maximum-intensity projections, and adjust brightness and contrast.

#### RNA detection and measurements

For quantification of single molecule RNA distribution relative to centrosomes, Nikon .nd2 files were first opened in Fiji, split into individual channels, and saved as .tif files using a custom macro. Raw images were then segmented using code adapted from the Allen Institute for Cell Science Cell Segmenter (Chen et al., 2018). Images were processed in batch using custom code written in Python and implemented using Jupyter notebooks. To minimize bias, we applied the same segmentation code to segment RNA objects under different biological conditions. Each segmented image was compared to the original image to validate accurate segmentation. RNA objects ≥ 50 pixels in segmented images were identified using the scikit-image tool label (van der Walt et al., 2014). Object features were then extracted using the regionprops tool from scikit-image. Extracted features included the raw image total pixel intensity, the object centroid coordinates, and the surface coordinates. These features were stored in a relational database using PostgreSQL.

For each image, the distances between centroid RNA coordinates and centroid centrosome coordinates was measured using the numpy vector normalization tool norm (van der Walt, Colbert, & Varoquaux, 2011). We then measured the distance between surface coordinates for a select number of RNA-centrosome pairs. Approximately 100 RNA objects were manually inspected to compare distances measured using centroid coordinates compared to distances measured using surface coordinates. The closest surface-to-surface distance for any given RNA corresponded to a centrosome in the top two closest centroid distance measurements. For this reason, the three closest centrosomes detected by centroid distance measurements were selected for surface measurements, to ensure that the closest centrosome was detected. This approach minimized processing time. The distance between the surface of each RNA object and its closest centrosome was recorded in the PostgreSQL database.

For single molecule normalization, we defined single molecules of RNA as RNA objects containing 50-100 pixels. These thresholds were selected based on the diffraction-limited 200 nm size of single RNA molecules detected by smFISH. For each RNA probe, we divided the integrated intensity of each RNA object by the averaged integrated intensity of all single RNA molecules, allowing an approximation of the number of RNA molecules per object, as previously described (Mueller et al., 2013). We then calculated the percentage of total RNA and percentage of total RNA in granules within a given distance from the centrosome (50 nm steps up to the pseudocell radius). We selected 10 μm and 4 μm as the pseudocell radius for NC 10 and NC 13, respectively, based on measuring the centrosome-to-centrosome distances from a set of representative images. The mean ± S.D. of the cumulative distributions were visualized using the Seaborn lineplot tool (Waskom et al., 2018).

#### Spindle morphology defects

Mitotic embryos imaged at 40x were examined for the following morphologies: bent spindles, multipolar or fused spindles, acentrosomal spindle poles, and defective centrosome separation. If any spindles within an embryo contained one of these phenotypes, the embryo was considered positive for a spindle morphology defect. Three independent biological replicates were performed for each genotype.

### Immunoblotting

Aged embryos were harvested, dechorionated in bleach, flash frozen in liquid nitrogen, and stored at −80 °C. 5-10 mg of frozen embryos were lysed with a 1 mL glass dounce homogenizer (Wheaton) in 100 μL lysis buffer (50 mM HEPES, 150 mM NaCl, 2.5 mM MgCl_2_, 0.1% Triton X-100, and 250 mM sucrose supplemented with 1x EDTA-free protease inhibitor cocktail (Roche, 04693159001), 1 μg/mL Pepstatin A (Sigma, P5318), 1 mM DTT (Sigma, 10197777001), and 2 mM RVC). 25 μL of 5x SDS loading dye was added to each lysate and samples were boiled for 10 min at 95 °C then resolved by SDS-PAGE gel and transferred to nitrocellulose membrane by wet transfer. Membranes were blocked for 1 hr at room temperature in a 5% dry milk solution diluted in TBST (Tris-based saline with 0.05% Tween-20), washed well with TBST, and incubated overnight at 4 °C with primary antibodies. After washing with TBST, membranes were incubated for 1 hr in the following secondary antibodies diluted 1:5000 in TBST, 5% milk: goat anti-mouse HRP (Thermo Fisher, 31430) and goat anti-rabbit HRP (Thermo Fisher, 31460). Membranes were washed well in TBST, and bands were visualized with Clarity ECL substrate (Bio-Rad, 1705061) on a Bio-Rad ChemiDoc imaging system.

Densitometry was measured using Fiji software using the ROI measure tool. For each sample, the ratio between the protein of interest and a loading control (e.g. β −Tub) was calculated. The mean relative expression and standard deviation were calculated and normalized to the mean of the biological control. Three independent biological replicates were processed on the same gel.

The following primary antibodies were used: rabbit anti-Cen (1:1000; gift from T. Megraw), mouse anti-FMRP (1:100; DSHB clone 5A11); mouse anti-β-Tub (1:1000; DSHB E7); and mouse anti-Actin (1:1000; DSHB clone JLA20).

### Immunoprecipitation

∼30 mg of frozen embryos were lysed with a glass dounce in 100 μL lysis buffer (50 mM HEPES pH 7.4, 150 mM NaCl, 2.5 mM MgCl_2_, 250 mM sucrose, 0.1% Triton X-100) supplemented with 1x protease inhibitor cocktail, 1 μg/mL Pepstatin A, 1 mM DTT, 1U/μL RNase Inhibitor (New England Biolabs, M0314S), and 2 mM RVC. Lysates were cleared by centrifugation, and the supernatant was pre-cleared in 25 μL of washed Protein A/G magnetic agarose beads (Pierce, 88802), or blocked magnetic beads (Chromotek, bmp-20) for GFP-Trap of FMRP, to reduce non-specific binding. 0.1-volumes of pre-cleared lysates were reserved as input, while the remainder was immunoprecipitated for 2 hr at 4^°^C in the following antibodies: rabbit anti-GFP (Invitrogen, A-11122), rabbit anti-Cen, or no antibodies as a control, then transferred to 25 μL washed Protein A/G magnetic agarose beads for immunoprecipitation for 2 hr. GFP-Trap magnetic agarose beads (Chromotek, gtma-10) were used for FMRP. Beads were then washed well in IP buffer (lysis buffer with 8 U/mL RNase Out and 0.4 mM RVC) then resuspended in 100 μL IP buffer. 50 μL of the beads (20% of volume for GFP-Trap) were analyzed for protein content by SDS-PAGE as described above. RNA was extracted from the other 50 μL of beads (80% of volume for GFP-Trap) using TRI Reagent (Sigma, T9424) and then treated with TURBO DNase (Thermo Fisher, AM2238) prior to RT-PCR.

cDNA was synthesized from 500 ng of RNA using Superscript IV Reverse Transcriptase (Thermo Fisher, 18091050) according to the manufacturer’s protocol with (RT+) or without (RT-) reverse transcriptase. DNA was amplified by PCR using Phusion High Fidelity DNA Polymerase (New England Biolabs, M0530L).

The following primers were used:

*cen* forward 5’-TAACCGCAGACGGACAAC-3’

*cen* reverse: 5’-GAATGCCCTATGGCTAGAAT-3’

*gapdh* forward: 5’-CACCCATTCGTCTGTGTTCG-3’ *gapdh* reverse: 5’-CAACAGTGATTCCCGACCAG-3’ *fmr* forward: 5’-CATCGTTCGACGGAGTAACA-3’ *fmr* reverse: 5’-GGAGCTTGTTGTTGGCTGAT-3’

### qPCR

RNA was extracted from ∼ 5 mg of frozen embryos using TRI Reagent, treated with Ambion Turbo DNase (Thermo Fisher, AM2238) for 30 min at 37 °C, followed by phenol:chloroform extraction. On the same day, RNA concentrations were measured with a spectrophotometer, and cDNA was synthesized from 500 ng of RNA using the iScript kit according to the manufacturer’s protocol (Bio-Rad, 170-8891).

qPCR was performed on a Bio-Rad CFX96 Real-time system with iTaq Universal SYBR Green Supermix (Bio-Rad, 172-5121). Three biological samples were tested in triplicate using 96 well-plates (Bio-Rad, HSP9601). *cen* expression levels were normalized to Ribosomal protein L32 (*RP49)*.

The following primers were used:

*cen* forward: 5’-TGAGGATACGACGCTCTGTG-3’

*cen* reverse 5’-AAAGTACCCCCGGTAACACC-3’, amplicon 78 bp;

*RP49* forward 5’-CATACAGGCCCAAGATCGTG-3’

*RP49* reverse 5’-ACAGCTTAGCATATCGATCCG-3’, amplicon 75 bp.

### Statistical Analysis

Data were plotted and statistical analysis was performed using Microsoft Excel and GraphPad Prism software. To calculate significance, the distribution normality was first assessed with a D’Agnostino and Pearson normality test. Data were then analyzed by Student’s two-tailed t-test, ANOVA, or the appropriate nonparametric tests and are displayed as mean ± SD. Data shown are representative results from at least two independent experiments, as indicated in the figure legends.

## Acknowledgements

We received gifts of reagents from Drs. Liz Gavis, Nasser Rusan, Tim Megraw, Greg Rogers, Ruth Lehmann, Mani Ramaswami, Martine Simonelig, Akira Nakamura, and Tom Jongens. We are grateful to Lauren Lym and Jina Lee for assistance with timed embryo collections and hatch rate analysis, respectively. We are indebted to Drs. Liz Gavis, Nasser Rusan, and members of the Lerit lab for helpful discussions and critical reading of this manuscript. Stocks obtained from the Bloomington *Drosophila* Stock Center (NIH grant P40OD018537); antibodies from the Developmental Studies Hybridoma Bank, created by the NICHD of the NIH and maintained at the University of Iowa Department of Biology; and reagents from the *Drosophila* Genomics Resource Center (NIH grant 2P40OD010949) were all used in this study. This work was supported by NIH grants 5K12GM000680 and 1F32GM128407 to PVR, AHA grant 20POST35210023 to JF, and NIH grant 5K22HL126922 to DAL.

---

**Figure 1 Supplement 1.**
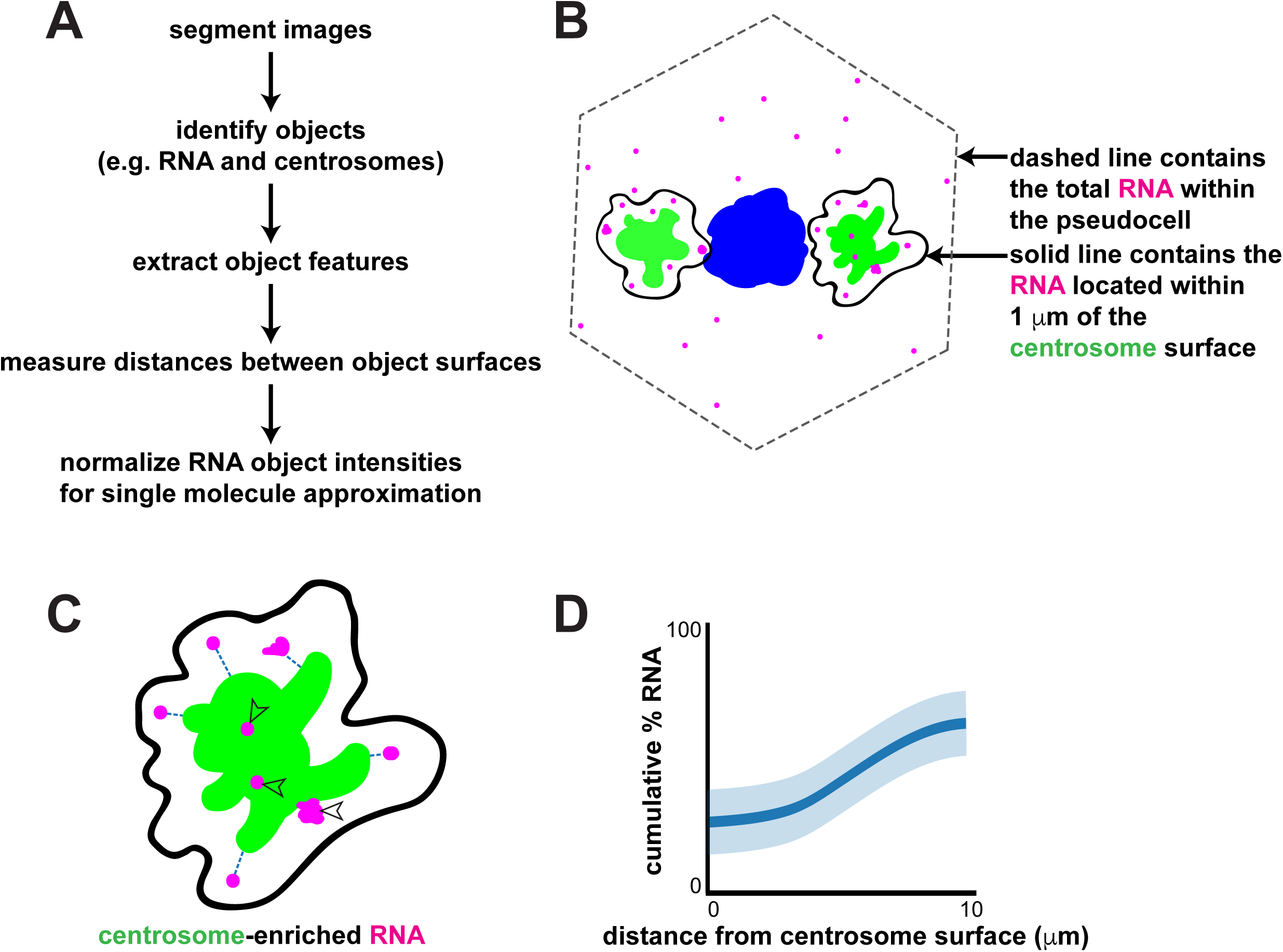
Schematic of image analysis for quantification of localized RNA. (A) Workflow used to quantify RNA distributions relative to centrosomes. (B) Cartoon shows the total RNA (magenta) within a syncytial *Drosophila* embryo pseudocell (dashed gray line) and RNA residing within 1 μm from a centrosome (green) surface (solid black lines). (C) The distances from the surfaces of each RNA object to the nearest centrosome were calculated (dashed lines). RNA objects that overlapped with centrosomes (open arrowheads) localized 0 μm from the centrosome. (D) Mock plot showing the cumulative distribution of RNA relative to distance from the centrosome, where 0 μm indicates RNA signals overlapping with centrosome signals. We define RNAs residing within 1 μm from the centrosome as centrosome-enriched (or pericentrosomal).

**Figure 1 Supplement 2.**
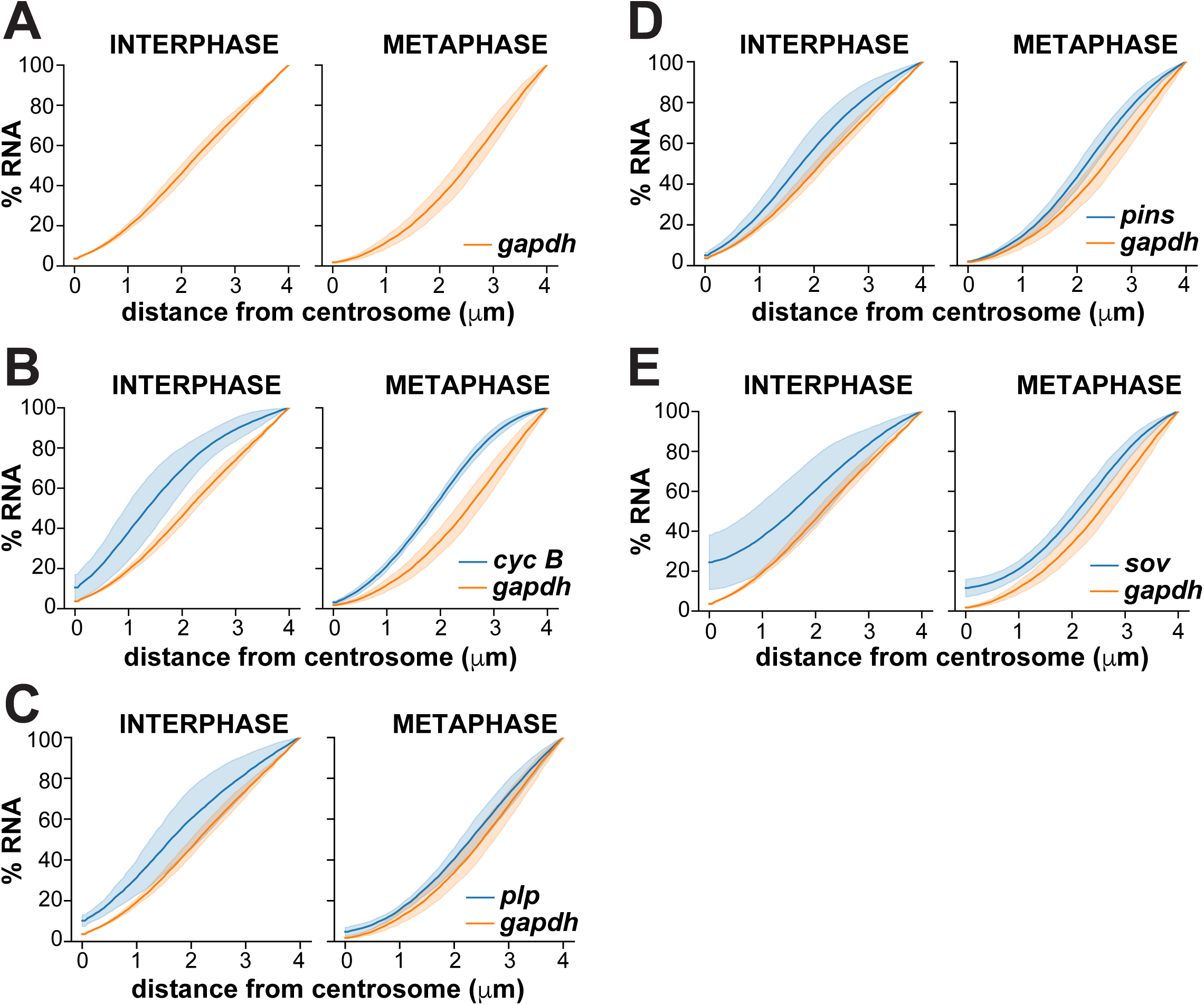
Cumulative distributions of centrosome-associated RNAs across the total cell volume. Graphs show the cumulative percentage of RNA as a function of distance from the centrosome surface as measured in NC 13 interphase or metaphase embryos. Data are plotted as mean ± S.D. (A) *gapdh*, (B) *cyc B*, (C) *plp*, (D) *pins*, and (E) *sov*. See Fig. 1 Supplemental Table 2 for details.

**Figure 1 Supplement 3.**
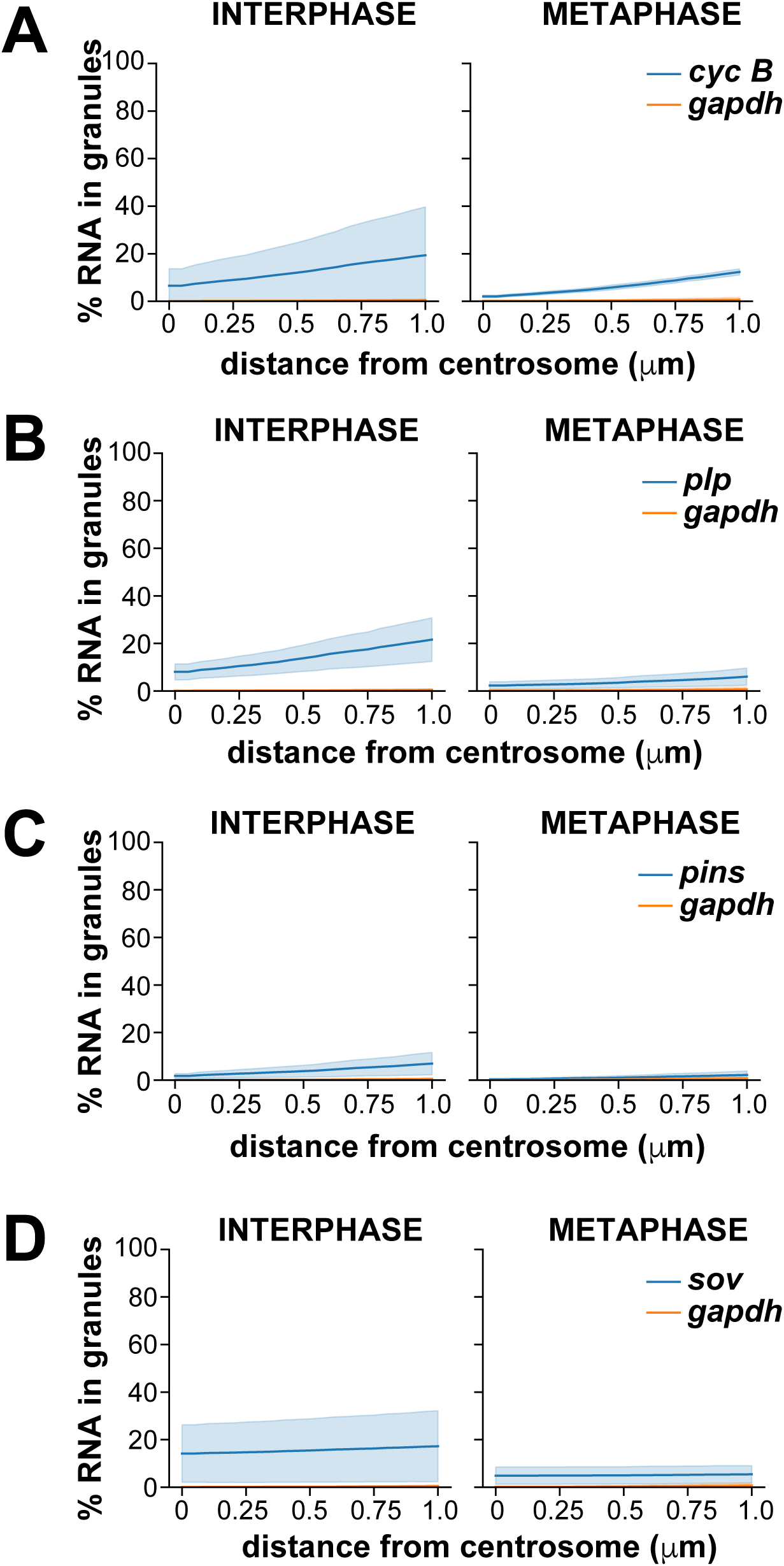
Cumulative distributions of granule-localized RNAs. Graphs show the cumulative percentage of RNA contained in granules (≥4 overlapping RNA objects) within 1 μm from a centrosome as measured in NC 13 interphase or metaphase embryos. Data are plotted as mean ± S.D. (A) *cyc B*, (B) *plp*, (C) *pins*, and (D) *sov*. See Fig. 1 Supplemental Table 2 for details.

**Figure 1 Supplement 4.**
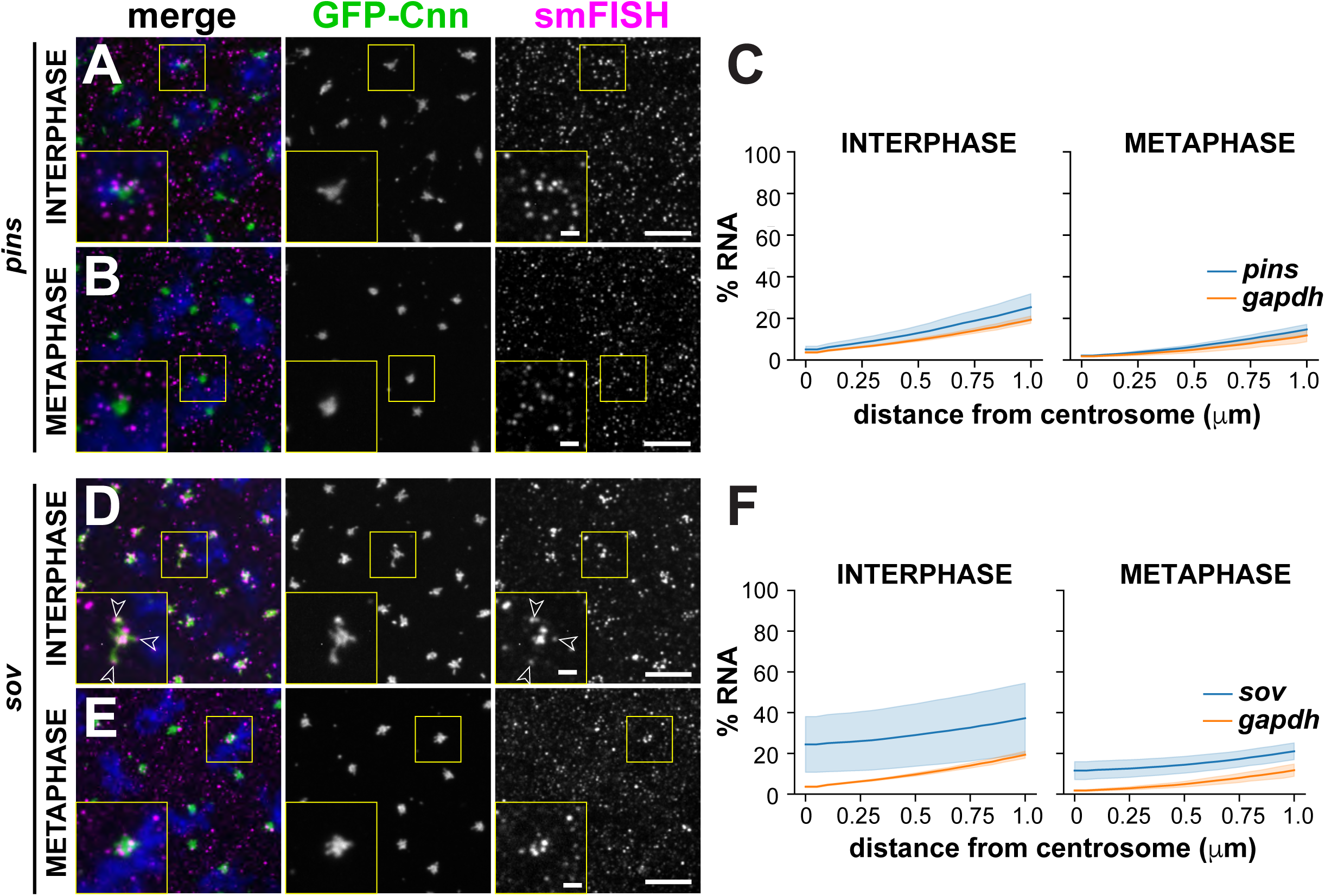
Localization of *pins* and *sov* mRNAs. Maximum intensity projections showing smFISH for *pins* or *sov* mRNAs (magenta) in interphase and metaphase NC 13 embryos expressing GFP-Cnn (green). Boxed regions are enlarged in the insets. Open arrowheads denote association of *sov* mRNA with centrosome flares. Quantification of the cumulative percentage of RNA located within 1 μm of the centrosome surface is shown to the right and plotted as mean ± S.D. (A–C) *pins* and (D–F) *sov*. See Fig. 1 Supplemental Table 2 for details. Scale bars: 5 μm and 1 μm (insets).

**Figure 2 Supplement 1.**
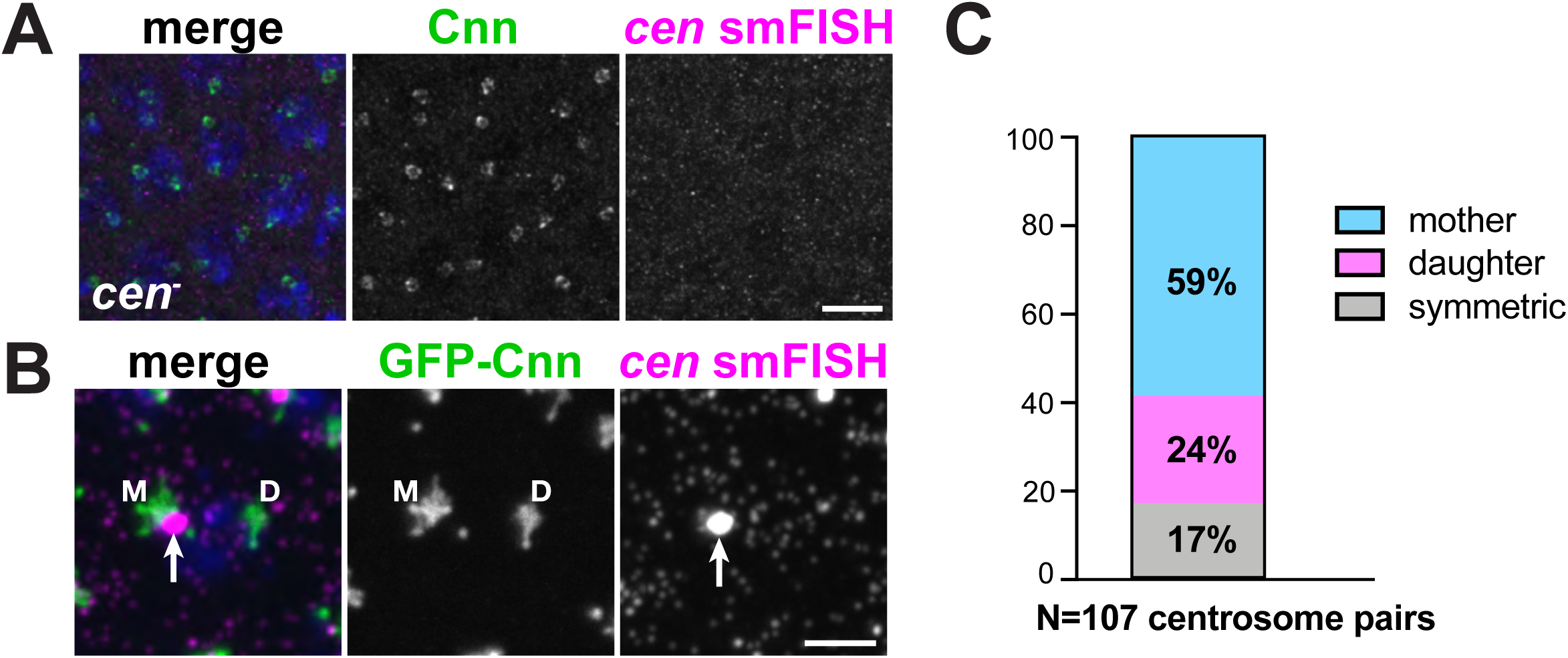
The *cen* granule preferentially localizes to the mother centrosome. (A and B) Maximum intensity projections showing *cen* smFISH (magenta) in NC 13 embryos relative to Cnn (green). (A) *cen* smFISH signals are not detected within null *cen* mutant embryos. Centrosomes are labeled with anti-Cnn antibodies. (B) An embryo expressing GFP-Cnn where the mother (M) and daughter (D) centrosomes are labeled. Arrows mark a *cen* granule localizing to the mother centrosome. (C) Quantification shows the frequency distribution of *cen* granule localization to the mother or daughter centrosome. N=107 centrosome pairs were measured from n=5 embryos. Scale bars: 10 μm (A) and (B) 2.5 μm.

**Figure 2 Supplement 2.**
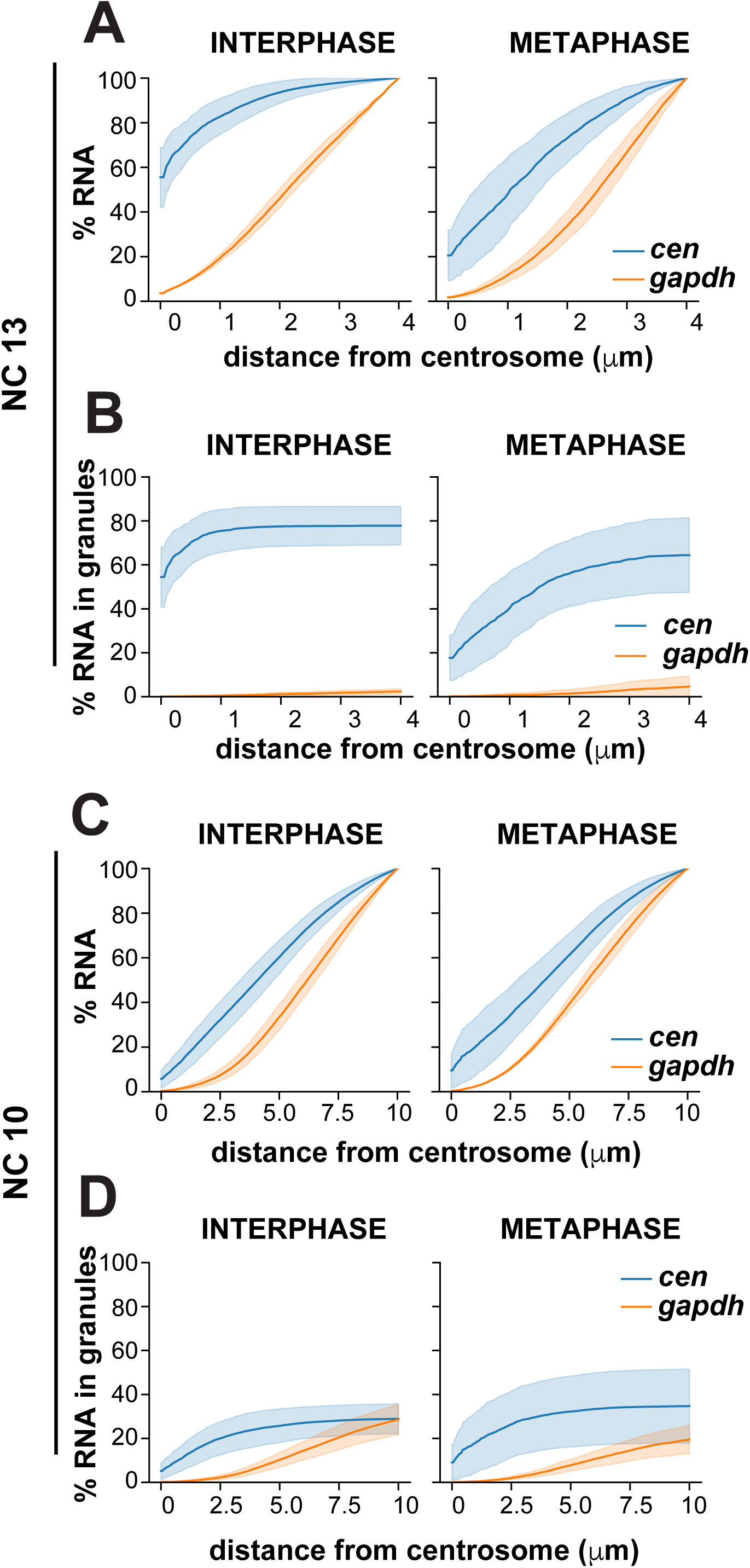
Cumulative distributions of *cen* RNA across the total cell volume. Graphs show the cumulative distributions of *cen* RNA (blue lines) relative to *gapdh* (orange lines) in interphase and metaphase embryos. (A and B) During interphase, NC 13 embryos show a majority of *cen* mRNA resides at centrosomes as a result of the accumulation of *cen* within pericentrosomal granules. (C and D) NC 10 embryos show more modest centrosomal enrichments of *cen* mRNA. Data are plotted as mean ± S.D. See Fig. 1 Supplemental Table 2 for details.

**Figure 3 Supplement 1.**
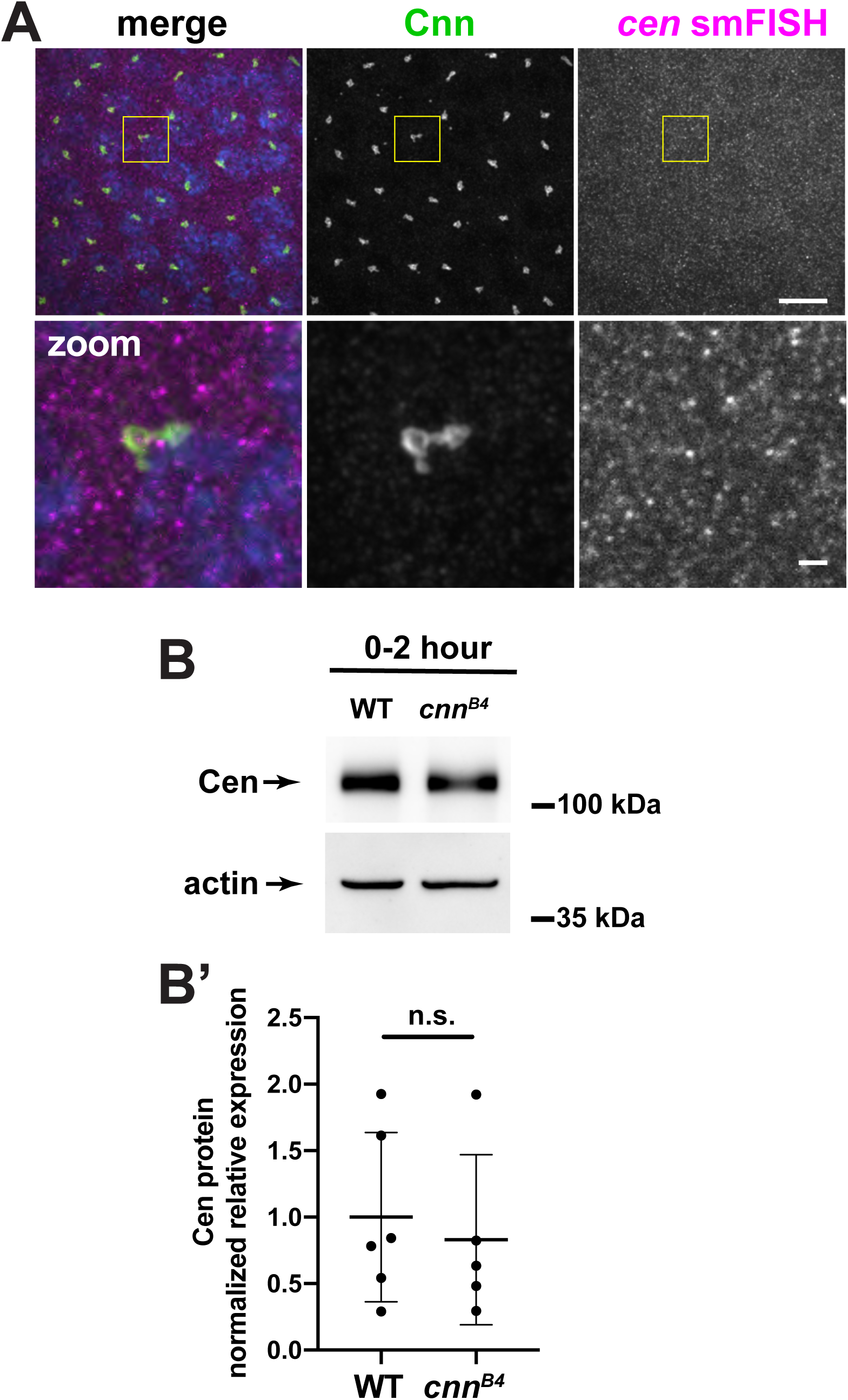
*cen* granule formation requires the centrosome scaffold. (A) Image shows immunofluorescence for Cnn (green) and *cen* smFISH (magenta) in an NC 12 *cnn*^*B4*^ embryo. Boxed region is enlarged below. Note the absence of large pericentrosomal *cen* granules. (B) Immunoblots show Cen protein content in 0-2 hour WT and *cnn*^*B4*^ lysates. Actin is used as a loading control. (B’) Graph shows the normalized expression levels of Cen. Each dot represents the levels of Cen normalized to the mean relative expression of the Actin load control. n.s. not significant (P=0.672) by unpaired t-test. Scale bars: 10 μm and 1 μm (insets).

**Figure 3 Supplement 2.**
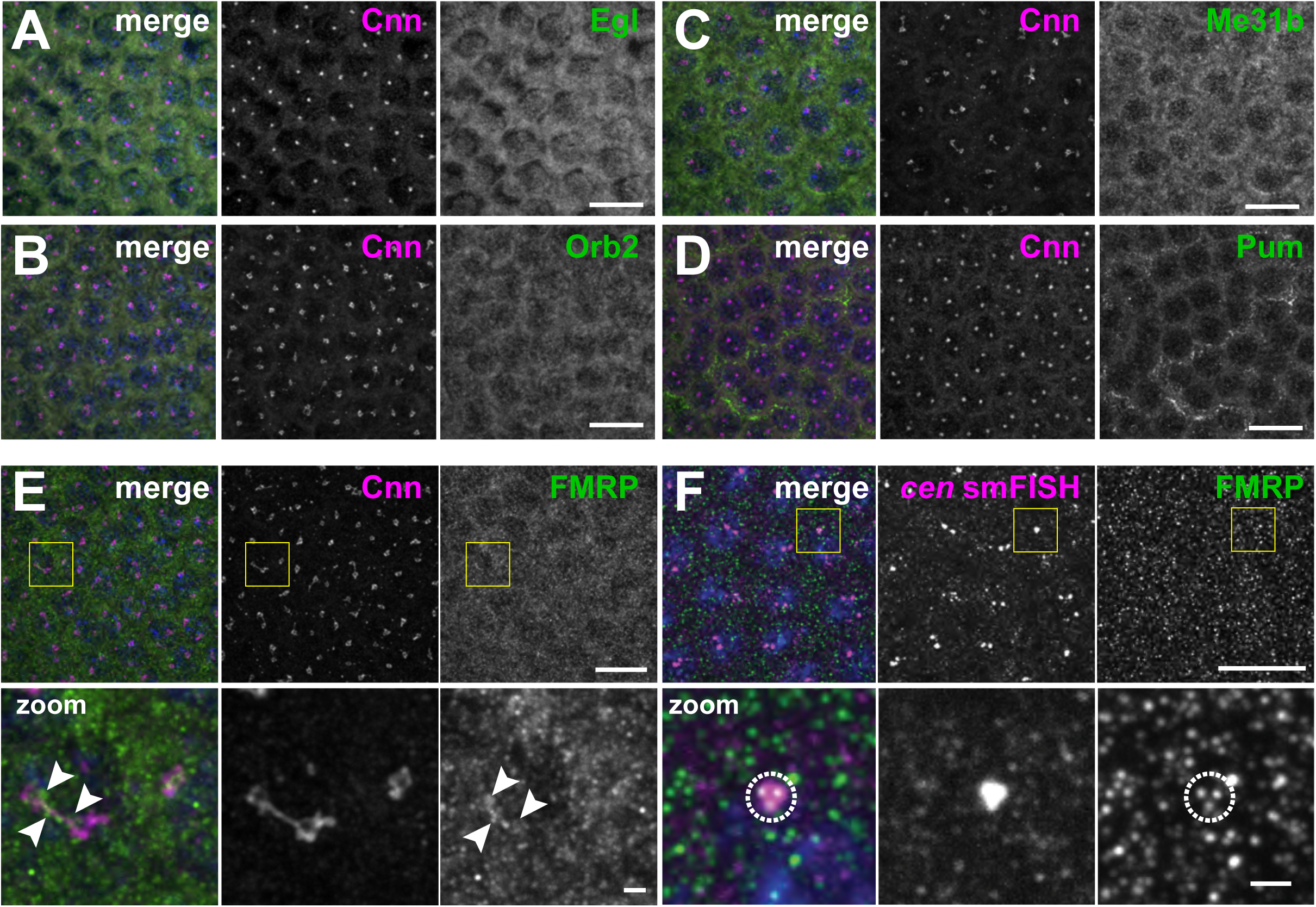
Candidate-based screen for centrosomal RNA-binding proteins. Images show interphase NC 12 embryos stained with Cnn (magenta) and antibodies for the indicated RNA-binding proteins (green): (A) Egl, (B) Orb2, (C) Me31B, (D) Pum, and (E) FMRP. Arrowheads shows FMRP overlapping with Cnn. (F) Immunofluorescence for FMRP was coupled with *cen* smFISH. Dashed circle marks FMRP puncta overlapping with *cen* RNA. Boxed regions are enlarged below. Scale bars: 10 μm and 2 μm (insets).

**Figure 4 Supplement 1.**
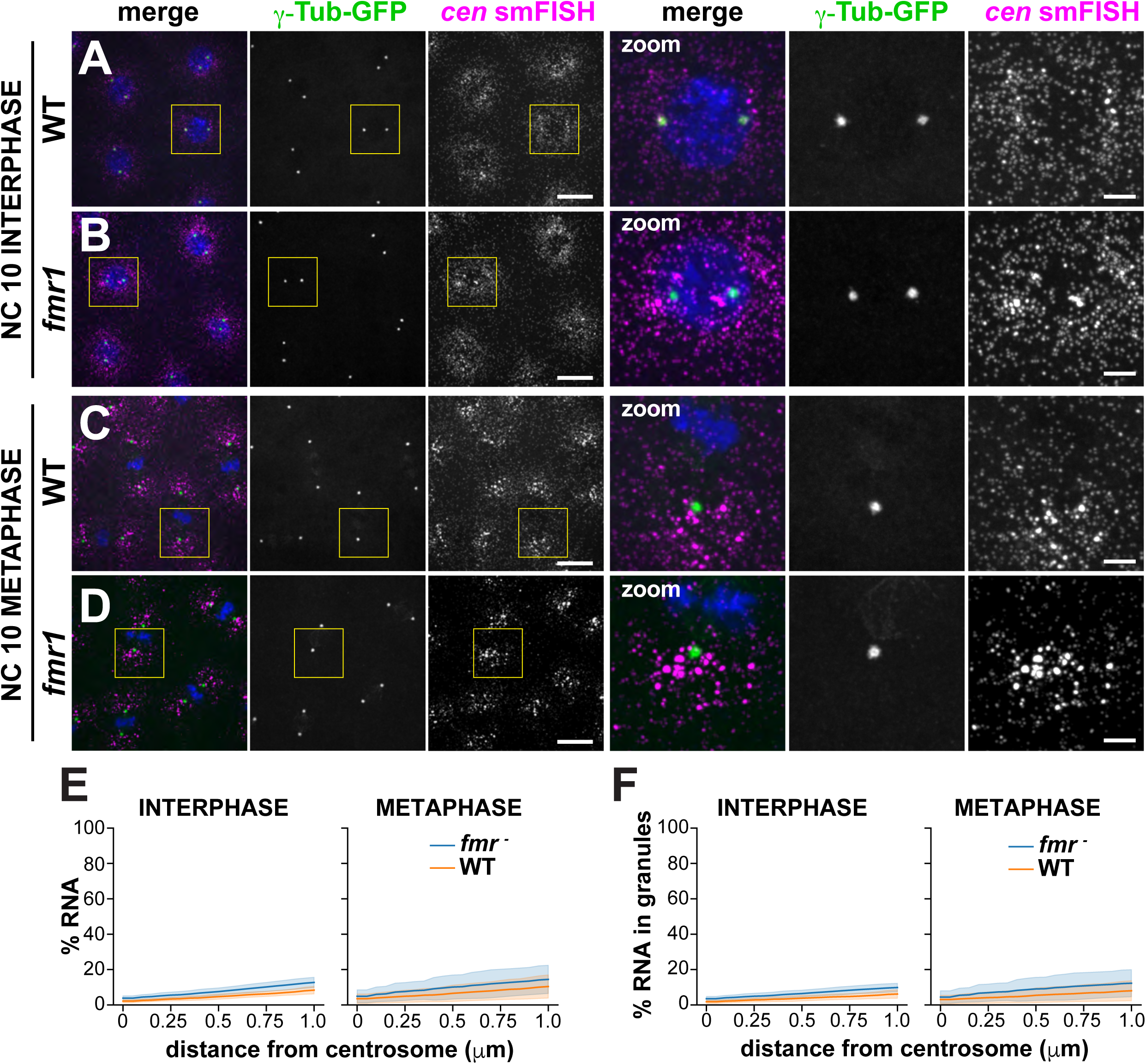
FMRP instructs the timing of *cen* RNA granule formation. Images show maximum intensity projections of WT or *fmr1* mutant NC 10 embryos expressing γTub-GFP and labeled with *cen* smFISH during (A and B) interphase or (C and D) mitosis. Boxed regions are enlarged to the right (zoom). (A) In interphase control embryos, *cen* is largely distributed as single molecules. (B) More *cen* granules are observed in interphase *fmr1* embryos. (C) In controls, *cen* granules form during mitosis. (F) In *fmr1* mutants, more *cen* is organized as granules. (E) Cumulative percentage of *cen* within 1 μm of the centrosome surface in WT (orange) or *fmr1* mutant (blue) embryos. (F) Cumulative percentage of *cen* within RNA granules up to 1 μm from the centrosome surface. Data are plotted as mean ± S.D. Note the similarity of the cumulative distribution plots in (E) and (F), indicating that the majority of the *cen* transcripts are contained within granules in both genotypes. See Fig. 4 Supplemental Table 1 for details. Scale bars are 10 μm and 2.5 μm (insets).

**Figure 4 Supplement 2.**
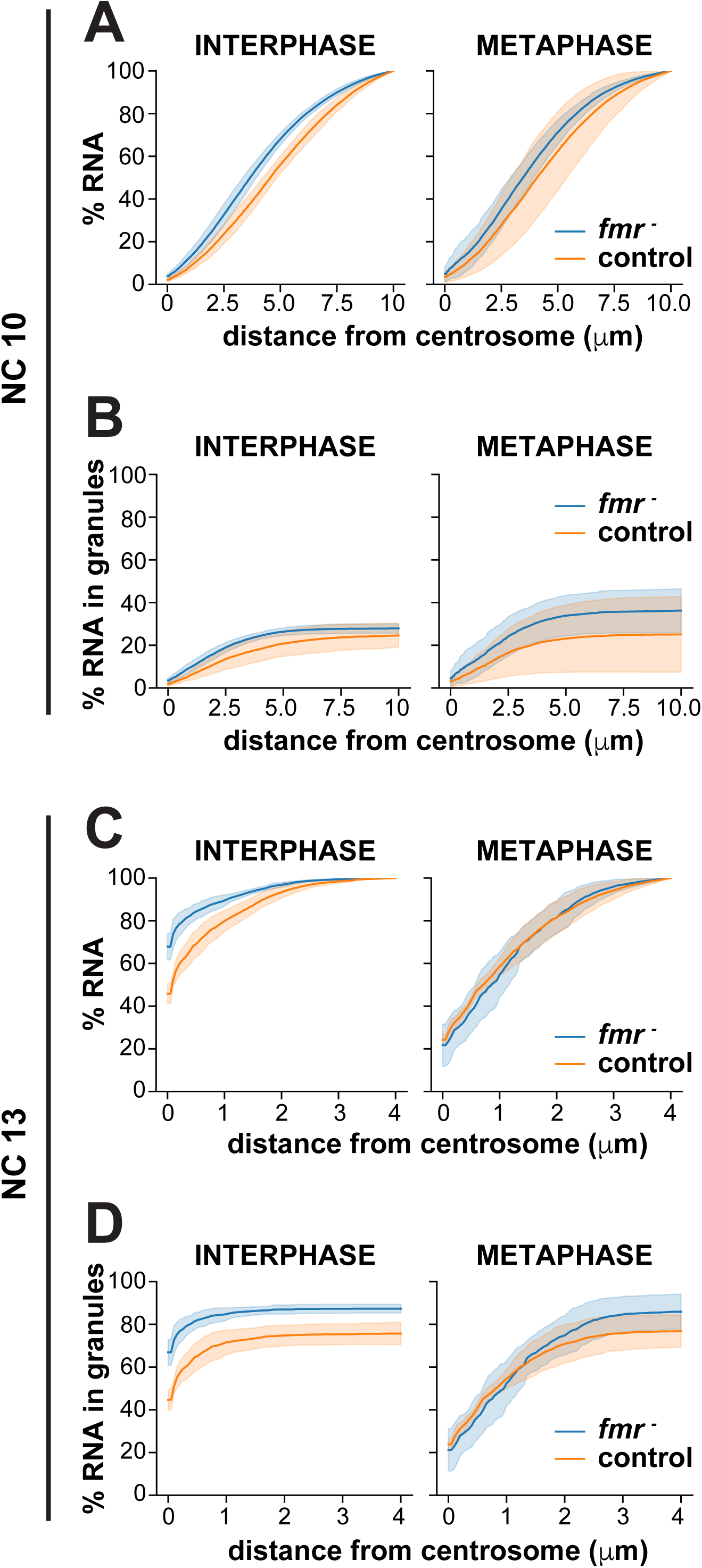
Cumulative distributions of *cen* RNA across the total cell volume in *fmr1* mutants. Graphs show the cumulative percentage of *cen* RNA as a function of distance from a centrosome surface, as measured in WT (orange lines) and *fmr1* mutant (blue lines) (A and B) NC 10 and (C and D) NC 13 interphase or metaphase embryos. Data are plotted as mean ± S.D. See Fig. 4 Supplemental Table 1 for details.

**Figure 6 Supplement 1.**
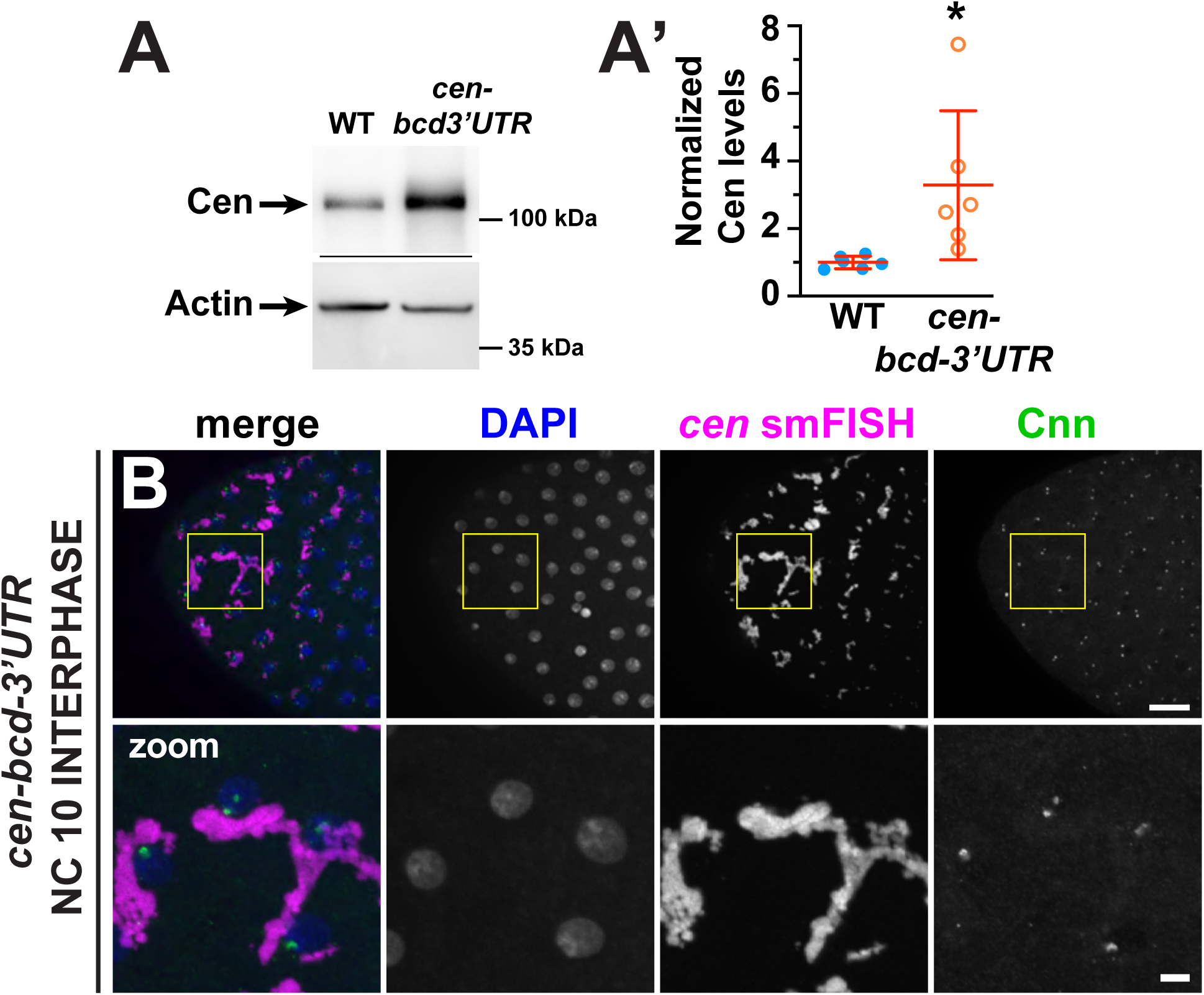
Cen protein expression in *cen-bcd-3’UTR* embryos. (A) Immunoblots show Cen protein content relative to the actin loading control from 1-3 hour embryonic extracts and are quantified in (A’). Levels of Cen were normalized to the mean WT levels of actin from N=3 independent biological replicates, each with n=2 technical replicates run on the same gel. (B) Maximum intensity projection of an early interphase NC 10 *cen-bcd-3’UTR* embryo. Note the formation of large-scale *cen*-containing RNPs adjacent to centrosomes. * P<0.05 by unpaired t-test. Scale bars: 10 μm and 2 μm (insets).

**Figure 1 Supplemental Table 1. Candidate centrosomal RNAs**. Genes documented to localize to centrosomes or spindle poles in (Lécuyer et al., 2007). Listed are the cDNAs used to generate traditional FISH probes in the Lécuyer screen (Column A) and the corresponding gene identifier information (Columns B and C). Additional information is annotated at http://fly-fish.ccbr.utoronto.ca/.

**Figure 1 Supplemental Table 2. Quantification of RNA localization to centrosomes**. For each mRNA analyzed (Column A), we documented the number of embryos (Column B), centrosomes (Column C), and RNA objects (Column D) quantified within NC 10 and 13 embryos in interphase versus metaphase. For each biological condition, we calculated the mean (Column E) and standard deviation (Column F) for the percentage of RNA within 1 μm of the centrosome surface per image. We used these data to calculate the mean fold-enrichment of mRNA relative to *gapdh* (Column G). We also calculated the mean (Column H) and standard deviation (Column I) percentage of RNA contained in granules containing 4 or more transcripts within 1 μm of the centrosome surface.

**Figure 1 Supplemental Table 3. smFISH probe sequences**. 5’ to 3’ smFISH probe sequences for *cen, cyc B, pins, plp, sov*, and *gapdh* are provided.

**Figure 4 Supplemental Table 1. Quantification of *cen* localization to centrosomes in WT and *fmr1* embryos**. For each given genotype (Column A), developmental stage (Column B), and cell cycle stage (Column C), we documented the number of embryos (Column D), centrosomes (Column E), and RNA objects (Column F) quantified. For each biological condition, we calculated the mean (Column G) and standard deviation (Column H) for the percentage of RNA overlapping with the centrosome surface per image. We used these data to calculate the mean fold enrichment of mRNA relative to the WT control (Column I). We also calculated the mean (Column J) and standard deviation (Column K) percentage of RNA contained in granules containing 4 or more transcripts overlapping with the centrosome surface. We repeated these same calculations for the volume within 1 μm of the centrosome surface (Columns M–Q).

**Table 1. *cen* overexpression increases embryonic lethality**. (A) Lethality rates in *fmr1* embryos and *fmr1* embryos that are hemizygous at the *cen* allele. (B) Lethality rates in *cen*-null embryos and embryos expressing the *cen-bcd-3’UTR* transgene in a *cen*-null background.

## Notes

Funding: This work was supported by NIH grants 5K12GM000680, 1F32GM128407 (PVR), and 5K22HL126922 (DAL) and an American Heart Association Postdoctoral Fellowship (20POST35210023) to JF.

### Competing Interest Statement

The authors have declared no competing interest.

